# Adaptive-like features of the γδ TCR couple chronic BTNL recognition to NK-like tissue immunity

**DOI:** 10.64898/2026.09.17.751481

**Authors:** Benjamin D McDonald, Hope D Anderson, Toufic Mayassi, Chhon Ling Sok, Veronica Locher, Magdalena Justyniarska, Megan E Borregard, Caroline Kaiser, Kristin Ladell, James E McLaren, David A Price, Narutoshi Hibino, Andrew S Koh, Jamie Rossjohn, Benjamin S Gully, Samantha J Riesenfeld, Bana Jabri

## Abstract

Intestinal Vγ4 γδ intraepithelial lymphocytes (IELs) persistently bind the constitutively expressed epithelial ligand BTNL3/8 through germline-encoded T cell receptor (TCR) determinants. While they resemble innate-like T cells such as NKT cells, unlike these populations, Vγ4 IELs encounter ligand only after thymic development. Moreover, unlike conventional αβ T cells, Vγ4 IELs sustain persistent physiological ligand engagement without becoming exhausted. Here, using biophysical, functional, and multimodal single-cell approaches, we show how Vγ4 IELs address this challenge. While germline-encoded TCR regions broadly mediate BTNL3 recognition, productive activation requires additional non-germline TCR features that license responsiveness to BTNL3/8 and enable local selection in the gut. Rather than driving exhaustion, BTNL3/8 reactivity directly promotes expression of an NK-like program marked by adaptor molecules that license innate-like signaling in healthy tissue. These findings reveal how combined innate and adaptive features of the γδTCR enable durable tissue specialization under conditions of persistent physiological ligand engagement.

## INTRODUCTION

Tissue-resident T cells and innate-like lymphocytes, such as NKT and MAIT cells, play essential roles in tissue protection and homeostasis. Vγ4 T cells that reside in the human intestine (Vγ4 IELs) represent a unique population that combines features of both T cell lineages. Like many innate-like T cells, Vγ4 IELs recognize their physiological ligand, BTNL3/8, through germline-encoded determinants within their TCRs^1–3^. However, unlike innate-like T cells that experience ligand-dependent programming during thymic development while they are undergoing lineage differentiation, Vγ4 IELs first encounter their constitutively expressed epithelial ligand, BTNL3/8, only after entering the gut as mature T cells^1,4^. How mature tissue-resident lymphocytes adapt to persistent physiological TCR engagement after tissue entry remains poorly understood.

In conventional T cells, including those that reside in the tissue, chronic TCR stimulation is classically associated with dysfunction or exhaustion rather than with the acquisition of innate-like effector properties^5^. However, in a subset of patients with active celiac disease, TCRαβ IELs instead acquire NK-like properties, including the expression of ITAM-bearing adaptor molecules that enable signaling pathways capable of bypassing TCR control over key effector functions^6^. A similar NK-like program has been reported in Vγ4 IELs in healthy tissue^7^. These observations raise the question of whether or how chronic TCR-ligand reactivity, in both inflammatory and homeostatic contexts, could instruct tissue-resident T cells to undergo this form of innate-like reprogramming. Since Vγ4 IELs have a defined ligand in healthy tissue, they make for a unique model for understanding the TCR requirements that couple chronic physiological ligand engagement to both tissue selection and durable functional specialization in mature T cells.

Recognition of BTNL3/8 has been proposed to be mediated by the hypervariable 4 (HV4) region of the Vγ4 chain^2,3^, suggesting that BTNL3/8 sensing may represent a broadly shared property of Vγ4 TCRs. However, most studies analyzing BTNL3/8 γδ T cell reactivity have examined Vγ4 IELs isolated from healthy intestinal epithelium, where BTNL3/8 is continuously expressed. In celiac disease, where BTNL3/8 expression is lost, we previously found that the proportion of Vγ4 IELs was significantly decreased, and that Vγ4 IEL TCRs that could be isolated from this setting were not reactive to BTNL3/8 in vitro^7^. These observations suggest that germline-encoded BTNL3/8 recognition alone is insufficient to drive the selection of Vγ4 γδ T cells within the intestinal epithelium. Moreover, γδ IELs in healthy tissue have been found to express an NK-like program that is lost along with BTNL3/8 expression in celiac disease and colorectal cancer, although it has not yet been determined if this program is directly linked to BTNL3/8 reactivity or other physiological cues.

This study addresses the fundamental, unresolved aspects of these previous findings. That is, if BTNL3/8 recognition is largely hardwired through germline TCR determinants, how are only a subset of Vγ4 IELs selected and durably specialized within the intestinal epithelium, while avoiding the dysfunction typically associated with persistent TCR engagement? More specifically, what TCR features enable physiological ligand binding to be translated into productive activation, tissue selection, and functional specialization?

To address these questions, we combined direct binding studies, cellular reactivity assays, and paired TCR single-cell transcriptomic analyses of human intestinal Vγ4 IELs. We show that BTNL3 binding and productive activation are mechanistically distinct processes. While germline-encoded determinants within the Vγ4 chain broadly mediate BTNL3 binding and induce partial signaling, productive responsiveness to BTNL3/8 requires additional adaptive-like non-germline determinants within both the γ and δ chains of the TCR. Strikingly, only Vγ4 IELs whose TCRs contain these adaptive-like determinants and are capable of full BTNL3/8 activation are selectively enriched within the intestinal epithelium. Moreover, we identify a core NK-like transcriptional program tightly associated with BTNL3/8 responsiveness, characterized by expression of the immunoreceptor tyrosine-based activation motif (ITAM)-bearing adaptor molecules FcεRIγ and DAP12, which, together with activating NK receptors, suggest the acquisition of signaling modules classically associated with NK-cell function. Together, these findings reveal that an innate-like tissue lymphocyte population relies on adaptive-like TCR features to couple chronic physiological ligand engagement to tissue selection and acquisition of a specialized NK-like signaling program, defining a distinct paradigm of tissue adaptation by mature T cells.

## RESULTS

### Tissue-specific BTNL3/8 expression drives selection of Vγ4 IELs

Previous studies of Vγ4 reactivity to BTNL3/8 have focused on γδ TCR sequences isolated from the IEL compartment^2–4,7^. We hypothesized that intestinal BTNL3/8 expression may bias the Vγ4 repertoire towards BTNL3/8 reactivity. To test this hypothesis and evaluate BTNL3/8 reactivity across a broader population of Vγ4 cells, we isolated γδ T cells from the peripheral blood and thymus—tissues classically devoid of BTNL3/8 expression—and compared their response to Vγ4 T cells isolated from the intestinal epithelium of the duodenum or colon of healthy donors^8,9^ (Methods). BTNL reactivity of γδ IELs is typically assessed by downregulation of surface TCR expression^2–4,7,10,11^. To evaluate BTNL3/8 reactivity, we co-cultured ex vivo Vγ4^+^ T cells with either HEK cells transduced to stably express BTNL3/8 (HEK-BTNL3+BTNL8, or in brief, HEK-BTNL3+8) or untransduced controls, and quantified reactivity as the relative percent TCR downregulation, normalized to negative controls (Methods). We found that Vγ4 T cells from blood exhibited significantly lower reactivity to BTNL3/8 compared to Vγ4 IELs (Figure 1A and 1B).

**Figure 1.**
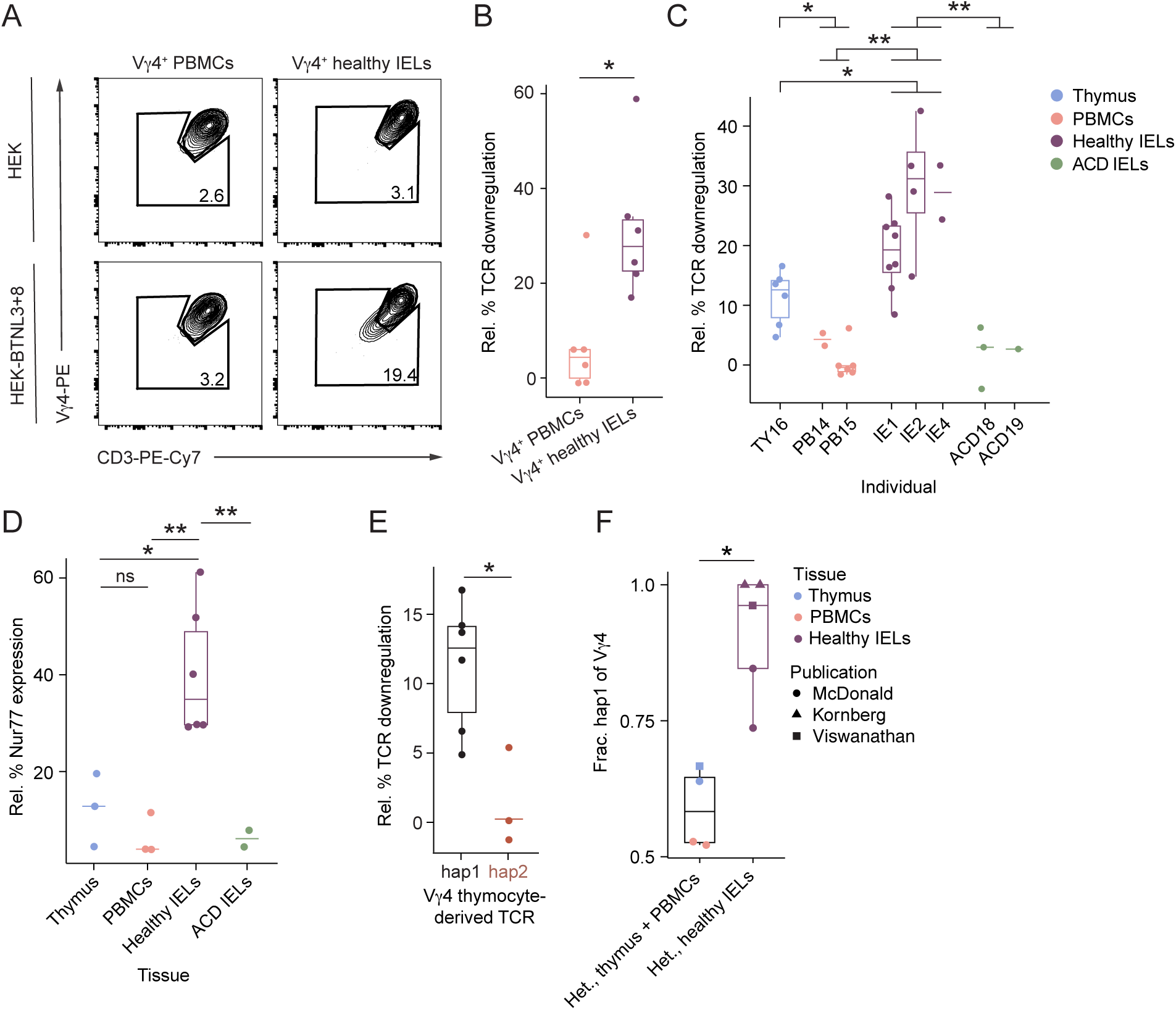
Tissue-specific BTNL3/8 expression drives selection of BTNL3/8-reactive IELs. (A) Representative flow plots display frequency of TCR downregulation (TCR^low^: Vγ4^low^CD3^low^) in Vγ4^+^ PBMCs (left) or Vγ4^+^ IELs from healthy individuals (right) cultured overnight with either HEK control cells (top) or HEK stimulator cells expressing BTNL3 and BTNL8 (“HEK-BTNL3+8”, bottom). (B) Boxplots show BTNL3/8 reactivity (relative % TCR downregulation; Methods) of Vγ4 cells isolated from tissues that do (healthy IELs, purple, *n*=6) and do not (PBMCs, pink, *n*=6) express BTNL3/8. Reactivity was measured by comparing co-cultures with HEK-BTNL3+8 to those with HEK cell controls. \**p*<0.05, two-tailed Student’s *t*-test. (C) Boxplots show BTNL3/8 reactivity (relative % TCR downregulation) of Jurkat cells expressing TCRs cloned from 8 individuals (x axis), as measured by comparing co-cultures with 100,000 HEK-BTNL3+8 cells to those with 100,000 HEK cells (controls) independently for each TCR (points; Methods). 32 distinct TCRs were cloned from individual samples of the thymus, PBMCs, and IELs from healthy individuals or those with active celiac disease (ACD) (color). \**p*<0.05, *\*\*p*<0.01; Holm-corrected p-values from one-tailed likelihood ratio tests (LRT) of mixed linear models accounting for individuals that do or do not include tissue labels (Methods). (D) Boxplots show an alternative measure of BTNL3/8 reactivity (relative % Nur77 expression; Methods) for a subset of tested TCRs (labeled and cultured as in C, but with 25,000 HEK-BTNL3+8 or 25,000 HEK cells). \*\**p*<0.01*, *p*<0.05, ns: *p ≥* 0.05; Holm-corrected p-values from two-tailed Student’s *t*-tests. (E) Boxplots show BTNL3/8 reactivity (relative % TCR downregulation) of Jurkat cells expressing γδ TCRs with distinct *TRGV4* haplotypes (x axis), derived from thymocytes of heterozygote TY16. \**p*<0.05, two-tailed Student’s *t*-test. (F) Boxplots show the fraction of cells using hap1 in all cells (using hap1, hap2, or hap3) isolated from tissues that do not (left; thymus, blue, and PBMCs, pink) or do (right; healthy IELs, purple) express BTNL3/8, from heterozygous individuals in two studies (our study, circle; Kornberg et al.^14^, triangle; Viswanathan et al.^15^, square). A single outlying blood patient (PB15) is excluded, but is visualized in Figure S2E and included in statistical test. *\*p*<0.05, LRT comparing beta regression model with and without a term considering gut residency (Methods; cell numbers per individual and haplotype in Table S1). Statistical analysis described per panel. For boxplots (shown if *n*>3), lower/middle/upper limits of boxes: first/second/third quartiles; whiskers extend to most outlying data point or 1.5x interquartile range, whichever is closest to median. Boxplots only shown for *n*>3; otherwise, horizontal bar indicates median. See also Figures S1–S2 and Table S1.

To validate these differences and control for potential cell- or tissue-intrinsic effects, including exposure to activating signals such as IL-15, which is enriched in the gut relative to the blood, we next assessed BTNL3/8 responsiveness using a Jurkat cell reporter system. Specifically, we performed paired single-cell γδ TCR-sequencing (scTCR-seq) and RNA-seq (scRNA-seq) experiments on γδ T cells isolated from the thymus (“Thymus”; 2 individuals), peripheral blood mononuclear cells (“PBMCs”; 3 individuals), IELs from healthy patients (“healthy IELs”, 11 individuals), and duodenal IELs from patients with active celiac disease (“ACD IELs”; 2 individuals; Table S1, Methods). A representative subset of the obtained TCR sequences (32/2,406 Vγ4 TCRs) were then cloned and expressed in Jurkat cells (Figure S1A–F), and BTNL3/8 reactivity was measured upon co-culture with HEK-BTNL3+8 cells. Stimulation with anti-CD3 was used as a positive control to confirm both cell-surface TCR expression and the capacity of each TCR to undergo downregulation following TCR crosslinking, thereby assessing its signaling competence and activation potential (Methods). For a subset of the tested TCRs, we validated reactivity results using Nur77 expression as an independent readout (Methods). While reactivity varied within and across donors, IEL-derived TCRs from healthy individuals exhibited significantly higher reactivity to BTNL3/8 than TCRs from other tissues or from active celiac disease patients (Figures 1C, 1D, S1G, and S1H; Table S2). Moreover, PBMC-derived TCRs were significantly less reactive than thymus-derived TCRs, which exhibited intermediate reactivity (Figures 1C and S1G). Overall, this suggests that BTNL3/8 expression in the gut selects for strongly reactive Vγ4 T cells.

Previous work identified haplotypic variants of *TRGV4* (Figure S2A; Vγ4 hap2 and Vγ4 hap3, versus the reference Vγ4 hap1 present in the IMGT database^12^) that account for up to 20% of *TRGV4* haplotypes in some populations^13^. These variants contain four non-conservative amino acid substitutions (and are therefore referred to as “haplotypes” rather than “alleles”). One substitution lies within the HV4 region, while the others are either solvent-exposed, located near the TCRδ chain, or positioned close to the TCRγ HV4 region, suggesting that they may influence ligand interactions (Figure S2B). Consistent with this, Corcoran et al. demonstrated that mutating a reactive hap1 TCR to hap3 reduced BTNL3/8 reactivity^13^. In our initial cloning assays, we tested only hap1 TCRs (Figure 1C and 1D).

To more deeply understand the impact of these variants on BTNL3/8 reactivity and selection for the IEL compartment, we first confirmed and extended previous findings. Specifically, we found that mutating a hap1 TCR to express either hap2 or hap3 reduced BTNL3/8 reactivity (Figure S2C–D). Our mutational experiments also identified a role for non-HV4 germline-encoded regions, in addition to the HV4 region previously found to play a role in BTNL3/8 reactivity (Figure S2C). To test whether *TRGV4* haplotype impacted BTNL3/8 reactivity in naturally occurring TCRs, we cloned TCRs containing hap1 or hap2 from the thymus of a hap1/hap2 heterozygous individual (TY16) and expressed them in Jurkat cells. The TCRs containing hap1 tended to be more reactive to BTNL3/8 than those with hap2 (Figure 1E).

To determine the functional relevance of Vγ4 polymorphisms *in vivo*, we then examined haplotype usage in the heterozygous, healthy individuals in our data, pooled with publicly available data from Kornberg et al.^14^ and Viswanathan et al.^15^ to increase sample size. We reasoned that because BTNL3/8 is not expressed in the thymus or the blood, the haplotype usage frequency in the thymus and PBMCs should be near 50-50, whereas, in the intestinal epithelium, where Vγ4 cells are selected for BTNL3/8 reactivity, haplotype usage should be skewed towards the more reactive haplotype hap1. Despite limitations in cohort size and IEL cell numbers, there was a striking preferential use of hap1 among Vγ4 IELs, consistent with a selective advantage of hap1 in the IEL compartment (Figures 1F and S2E, Table S1). In contrast, in Vγ4 cells from the thymus and PBMCs, hap1 usage was lower, except in a single hap1-biased outlying PBMC sample (PB15) whose Vγ4 TCRs had reduced clonotype diversity but were not reactive to BTNL3/8 (Figures S2F and 1C, Table S1), potentially suggesting hap1-biased selection by another ligand. Altogether, these results demonstrate substantial selective pressure in the IEL compartment for strongly BTNL3/8-reactive TCRs.

### CDR3 features and Vγ4–Vδ pairing cooperatively shape BTNL3/8 reactivity

The heightened BTNL3/8 reactivity we observed in the Vγ4 IELs, even among hap1 TCRs (Figures 1C–D), suggested that features beyond the germline-encoded *TRGV4* region—specifically, the CDR3 region—also play a role in shaping BTNL3/8 recognition. To isolate the impact of each chain, we studied each independently before examining the impact of γ–δ chain pairing on reactivity.

Focusing first on the γ chain, we analyzed the γ chain CDR3 repertoire of Vγ4 hap1 TCRs from IELs, where BTNL3/8 is expressed, and compared it with that from the thymus (in this study and a published dataset^16^), reasoning that the repertoires should be distinguishable if CDR3γ contributed to BTNL3/8 reactivity and selection into the IEL compartment. Adapting an approach from αβ TCR repertoire analysis^17^, we tested for enrichment of amino acid *k*-mer motifs in CDR3γ regions (Methods) and identified three motifs significantly enriched in TCRs from IELs relative to those from the thymus (Figures S3A–B; Table S3). Given our limited statistical power to detect enrichment of individual motifs, we also tested for signal in the distributions of motifs, which revealed a significant overall difference in the CDR3γ repertoire between IELs and the thymus (Figure 2A, Methods). While CDR3γ sequence length distributions were comparable between the two compartments (Figure S3C), there was a marked enrichment of JγP1 gene segment usage in IELs, compared to Vγ4 cells from the thymus (Figure 2B), further underscoring differences in the γ chain repertoire. Together, these findings suggest that both germline and non-germline features of the γ chain contribute to the selection of Vγ4 T cells into the IEL compartment.

**Figure 2.**
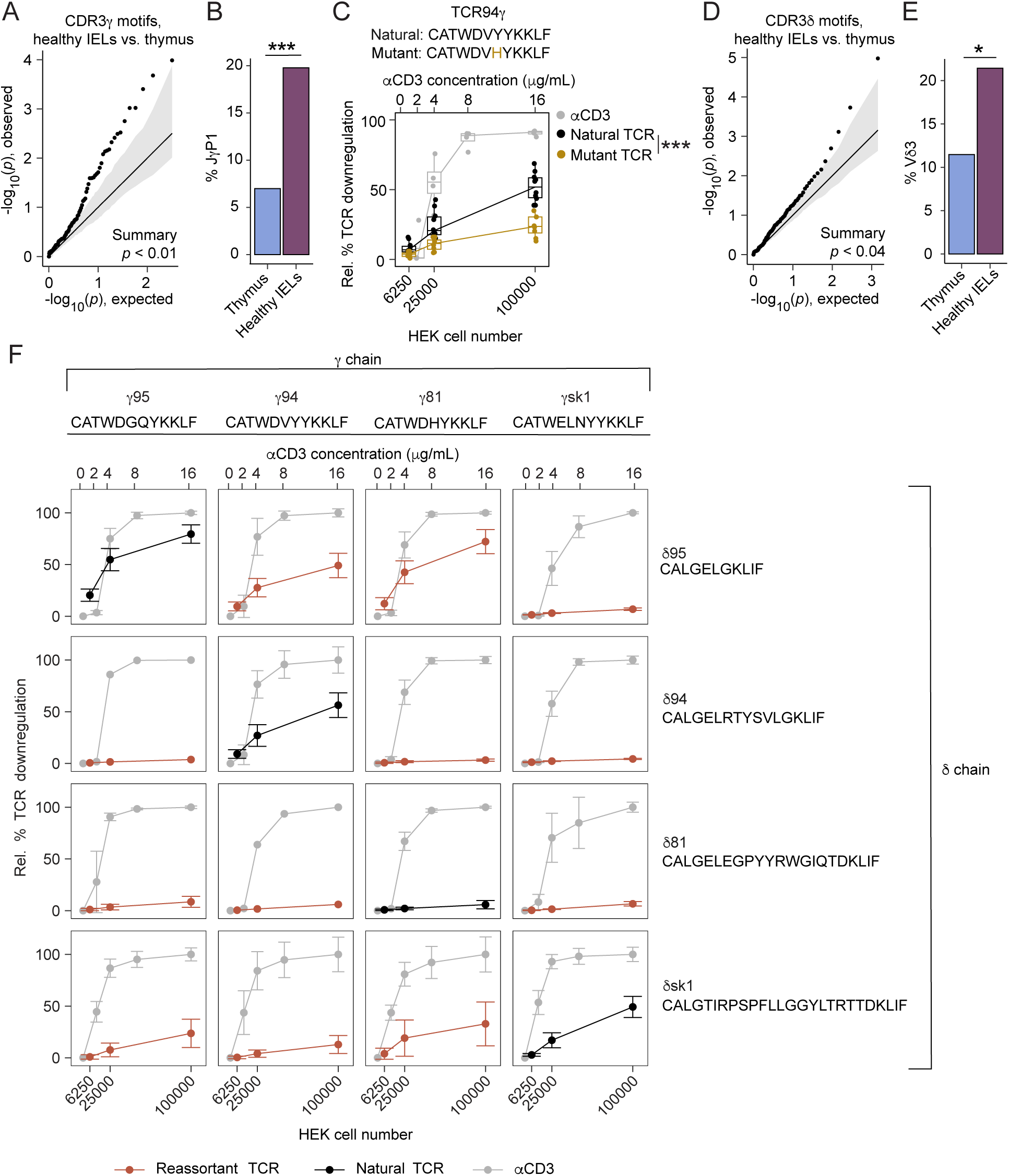
CDR3 features and Vγ-Vδ pairing cooperatively shape BTNL3/8 reactivity. (A) A Q-Q plot indicates overall differential enrichment in CDR3γ motifs in Vγ4 hap1 TCR clones from healthy IELs versus those from the thymus. The plot compares (1) quantiles of p-values from two-tailed Fisher’s exact tests (y axis, “observed”), each testing enrichment of a single CDR3γ motif, with (2) mean quantiles over 100 samples of an empirical null p-value distribution (x axis, “expected”), generated by performing similar enrichment tests for tissue-label permutations (IELs, thymocytes). Line: equal expected and observed distributions; gray area: 90% confidence interval, estimated from 5^th^ and 95^th^ percentiles of each quantile. The tail (*p*<0.05) of the observed distribution is significantly skewed toward smaller p-values (*p*<0.01, permutation test; Methods). (B) Bar plot shows the frequencies of Vγ4 hap1 TCR clones from the thymus and healthy IELs (x axis) that use the JγP1 gene segment. \*\*\**p*<0.001, two-tailed Fisher’s exact test, p-value is Holm-corrected for testing of all possible Jγ gene segments. (C) *Top*: The natural and mutant CDR3γ AA sequences of TCR94 are displayed, with the pointwise AA swap in the CDR3γ highlighted (goldenrod). *Bottom*: boxplots show BNTL3/8 reactivity (relative % TCR downregulation) of Jurkat lines expressing either the natural (black) or CDR3γ mutant (goldenrod), as determined by co-culturing with various numbers (bottom x axis) of either HEK control or HEK-BTNL3+8 stimulator cells. The relative % TCR downregulation for the mutant TCR stimulated with various concentrations of anti-CD3 (top x axis) is also shown (gray). \*\*\**p*<0.001, one-tailed LRT of linear models using minimum and maximum HEK cell numbers as independent variables, with or without TCR labels (Methods). Lower/middle/upper limits of boxes: first/second/third quartiles; whiskers extend to most outlying data point or 1.5x interquartile range, whichever is closest to median. (D) A Q-Q plot (analogous to A) indicates overall differential enrichment in CDR3δ motifs in Vγ4 hap1 TCR clones from healthy IELs versus those from the thymus. The tail (*p*<0.05) of the observed distribution is significantly skewed toward smaller p-values, compared with that of the null (*p*<0.04, permutation test, as in A). (E) Bar plot (analogous to B), for the Vδ3 gene segment. \**p*<0.05, two-tailed Fisher’s exact test, p-value is Holm-corrected for testing of all possible Vδ gene segments. (F) Plots show BTNL3/8 reactivity (relative % TCR downregulation) for Jurkat lines expressing naturally occurring TCRs (4 diagonal panels, black lines) or reassortant TCRs (12 off-diagonal panels, red lines) generated by pairing each natural TCRγ (column) with each natural TCRδ (row). Reactivity was determined by co-culturing each Jurkat line with variable numbers (bottom x axis) of either HEK control or HEK-BTNL3+8 stimulator cells, or different concentrations (top x-axis) of plate-bound anti-CD3. The relative % TCR downregulation at various concentrations of anti-CD3 (top x axis) is also shown (gray). 1–25 experiments were performed for each combination of parameters; points: mean; error bars: standard deviation. Statistical analysis described per panel. For boxplots, lower/upper limits of boxes: first/third quartiles; middle box line: median; whiskers extend to most outlying data point or 1.5x interquartile range, whichever is closest to median. Boxplots only shown for *n*>3; otherwise, horizontal bar indicates median. See also Figure S3 and Table S2.

To directly assess the contribution of the CDR3γ to BTNL3/8 reactivity, we performed a targeted mutagenesis experiment on TCR94, a previously characterized BTNL3/8-reactive TCR isolated from the duodenal epithelium of a healthy individual^7^. We substituted a histidine with a tyrosine in the CDR3γ (Figure 2C). The mutant TCR was expressed in Jurkat cells, where we confirmed proper surface expression and responsiveness to anti-CD3 stimulation, as shown by TCR downregulation (Figure 2C, light gray line). This single-residue change, which introduced a recurrent motif observed in celiac disease patients^7^, led to a significant reduction in BTNL3/8 reactivity (Figure 2C), demonstrating that even subtle differences in the CDR3γ can critically alter TCR responsiveness to BTNL3/8.

We next applied a similar approach to assess whether δ chain traits distinguish IEL and thymus repertoires of Vγ4 hap1 TCRs. As with the CDR3γ, CDR3δ motifs in Vγ4 hap1 TCRs differed significantly overall between IELs and the thymus (Figure 2D), with significant IEL enrichment detected for only a few individual motifs after correcting for multiple hypothesis testing (Figures S3D–E, Table S3). We also found that IEL-derived Vγ4 hap1 TCRs more frequently used Vδ3 gene segments (Figure 2E, Table S3) and exhibited slightly shorter CDR3δ regions than their thymic counterparts (Figure S3F). Together, these findings suggest that both germline and non-germline features of the δ chain contribute to the selection of Vγ4 T cells into the IEL compartment.

To directly assess the impact of the CDR3δ sequence on BTNL3/8 reactivity, we created a mutant of BTNL3/8-reactive TCR94 by inserting 5 glycines in the CDR3 (Figure S3G). We expressed the mutant TCR in Jurkat cells, verified its expression on the cell surface, and confirmed that it was reactive to anti-CD3 stimulation (Figure S3G). Jurkat cells expressing the CDR3δ mutant TCR94 showed reduced BTNL3/8 reactivity, relative to those with natural TCR94, upon co-culture with HEK-BTNL3+8 cells (Figure S3G). While these results are consistent with the implication of our computational analysis that the CDR3δ influences BTNL3/8 reactivity, and more specifically, that shorter CDR3δ may be more likely to react to BTNL3/8, we cannot disentangle the effects of manipulating CDR3δ length from strongly altering CDR3δ conformation in this assay. Regardless, taken together, the results expose a previously unknown role of the TCRδ chain in influencing Vγ4 BTNL3/8 reactivity.

As our data thus far independently implicated both γ and δ chains in influencing reactivity, we next asked whether BTNL3/8 recognition requires specific γδ pairings, or if individual, reactive γ or δ chains can drive reactivity regardless of their pairing partner. To specifically investigate the combined effects of CDR3γ and CDR3δ, we generated Jurkat cell lines expressing four previously published Vγ4 TCRs, referred to as “natural TCRs” (three reactive: 95, 94, sk1; one minimally reactive: 81; Table S2), whose established BTNL3/8 reactivity was validated in our TCR downregulation assay (Figure 2F, Methods)^3,7^. We then generated Jurkat lines expressing 12 “reassortant TCRs” by pairing each of the four natural TCRγ chains with each of the four natural TCRδ chains in all possible combinations. All TCRs were successfully expressed at the cell surface and were effectively downregulated by anti-CD3 stimulation (Figure 2F, Methods). BTNL3/8 reactivity assays of all TCRs showed that no single γ or δ chains was uniformly dominant. While naturally occurring TCR95 (γ95 + δ95) was strongly reactive, the pairing of γ95 with certain δ chains (δ94, δ81) abrogated reactivity. Analogously, pairing δ95 with a different γ chain (γsk1) also reduced BTNL3/8 reactivity (Figure 2F). Similar effects were observed across the combinatorial experiment (e.g., γ94 + δ94 vs. γ94 + δsk1 or γsk1 + δ94). Inversely, the low reactivity of naturally occurring TCR81 (γ81 + δ81) was increased dramatically by pairing γ81 with δ95 instead. Altogether, our data are consistent with a complex, multifaceted model of γδTCR-BTNL3/8 interaction, with key roles for both germline-encoded and non-germline-encoded amino acids of the TCRγ and δ chains, as well as the chain pairing, in influencing BTNL3/8 reactivity.

### Molecular binding and cellular reactivity of Vγ4 T cells are dissociated

To determine whether differences in BTNL3/8 reactivity reflected differences in BTNL3 binding, we performed experiments using Vγ4 TCRs with distinct functional profiles: the BTNL3/8-reactive TCRs 95 and 94, isolated from IELs in healthy tissue, and the minimally reactive TCR81, derived from the duodenal IEL compartment of a patient with active celiac disease (Table S2)^7^. A TCR downregulation assay (Figure 3A) and an NFAT-driven luciferase reporter assay (Figure 3B) confirmed that, in the presence of HEK-BTNL3+8 cells, TCRs 95 and 94 displayed significantly higher reactivity than TCR81, which was not explained by variations in TCR sensitivity to anti-CD3 stimulation (Figure S4A).

**Figure 3.**
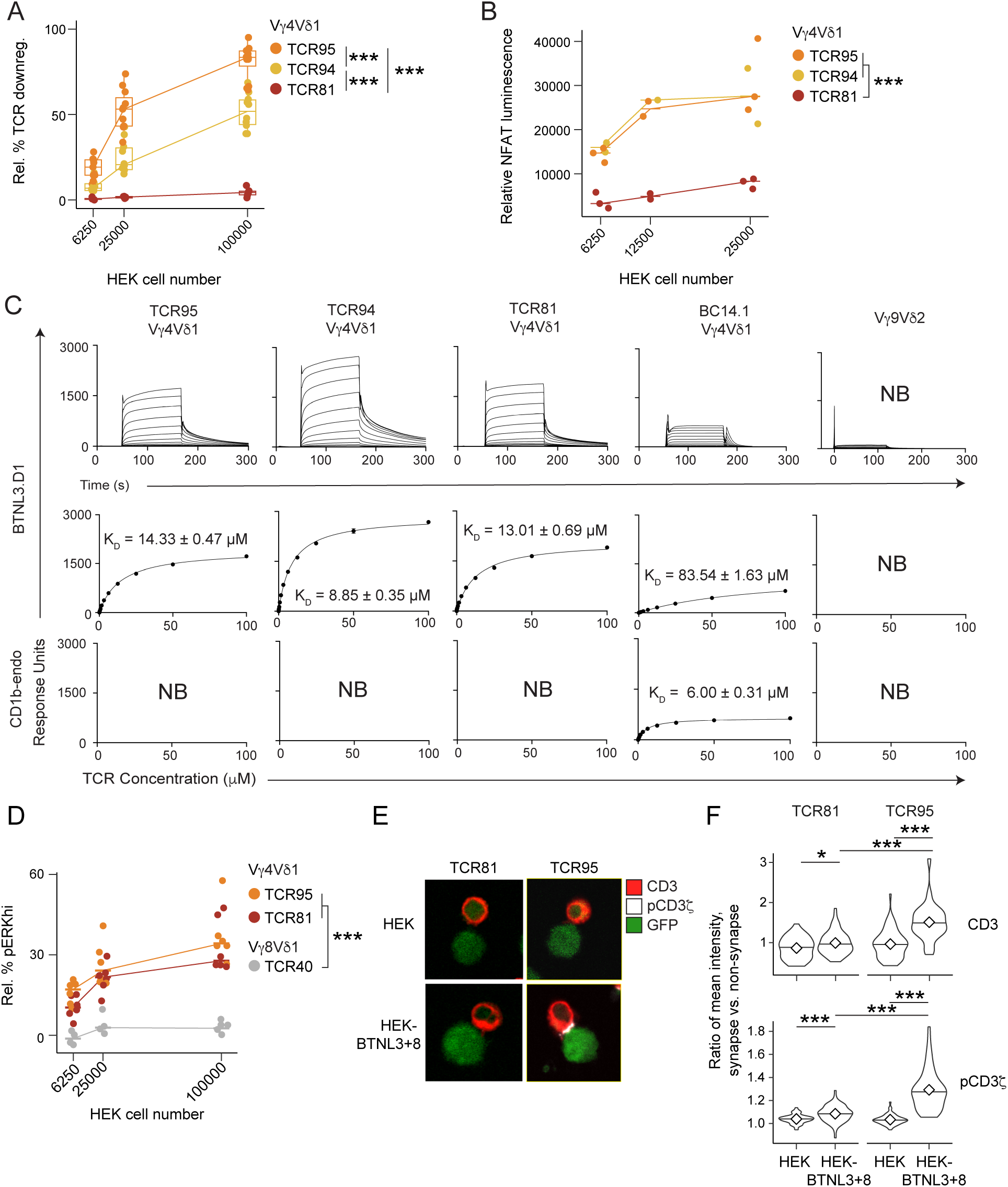
Molecular binding and cellular reactivity of Vγ4 T cells are dissociated. (A) –(B) Plots show BTNL3/8 reactivity via two different metrics: relative TCR downregulation (A, y axis) and batch-regressed NFAT-luciferase driven luminescence (B, y axis; Methods) for Jurkat lines expressing BTNL3/8-reactive TCRs (TCR95, orange; and TCR94, yellow) or a minimally BTNL3/8-reactive TCR (TCR81, red), each cultured as in Figure 1C, but with variable numbers of either HEK control or HEK-BTNL3+8 stimulator cells (x axis). For each parameter combination, *n*=9–14 (A) and *n*=2–3 (B). \*\*\**p*<0.001, Holm-corrected p-values from one-tailed LRTs of linear models using minimum and maximum HEK cell numbers as independent variables, with or without TCR labels (Methods). (C) Representative surface plasmon resonance (SPR) sensorgrams (top) and derived equilibrium binding affinity curves (bottom) are shown for naturally occurring γδ TCRs (TCR95, TCR94, and TCR81), a Vγ4 CD1b-reactive TCR (BC14.1), and a control Vγ9Vδ2 TCR. Each TCR was serially diluted from 100 to 0 µM and tested for binding with immobilized BTNL3 and CD1b at a surface density of ∼2000 response units. Sensorgrams from three independent experiments, with two replicates, were used to determine the dissociation constants (K_D_) ± SEM from a one-site specific binding model. Equilibrium binding curves show mean binding ± SEM for replicates or no binding (NB) where undetected. (D) Plot shows relative % pERK^hi^ (Methods) for Jurkat lines expressing a BTNL3/8-reactive Vγ4 TCR (TCR95, orange, as in A), a minimally BTNL3/8-reactive Vγ4 TCR (TCR81, red, as in A), and a BTNL3/8-non-reactive Vγ8 TCR (TCR40, gray), each cultured as in Figure 1C, but with variable numbers of either HEK control or HEK-BTNL3+8 stimulator cells (x axis). Horizontal bars: median; for each parameter combination, *n*=4–6. \**p*<0.001, Holm-corrected p-values from one-tailed LRTs of linear models using minimum and maximum HEK cell numbers as independent variables, with or without TCR labels (Methods). (E)–(F) Representative images (E) and summary quantification (F) of CD3 (red) and pCD3ζ (white) staining of Jurkat lines TCR81 (left) and TCR95 (right), co-cultured with either HEK control or HEK-BTNL3+8 stimulator cells (green) (rows, E; x axis, F). For HEK-Jurkat pairs, ROIs were defined at the immunological synapse and distal membrane regions. Mean fluorescence intensities were extracted from 50 pairs per condition and used to compute synapse vs. non synapse ratios. \**p*<0.05, \*\*\**p*<0.001, Holm-corrected p-values from two-tailed Student’s *t*-tests. Statistical analysis described per panel. Nested black brackets indicate statistical significance for each pairwise comparison. For boxplots (in A, shown if *n*>3), lower/middle/upper limits of boxes: first/second/third quartiles; whiskers extend to most outlying data point or 1.5x interquartile range, whichever is closest to median. For *n*≤3, horizontal bar: median. For violin plots (in F), diamond: median; horizontal black bands: 25th and 75th percentiles. See also Figures S4–S5.

We then used surface plasmon resonance (SPR) to assess the binding of soluble recombinant γδ TCRs to recombinantly expressed and immobilized IgV domains of BTNL3 and BTNL8. Despite marked differences in cellular activation among TCR95, TCR94, and TCR81, all three Vγ4 TCRs bound the BTNL3 IgV domain with affinities consistent with that of the Vγ4Vδ1 TCR BC14.1 to its known ligand CD1b^18^ (Figure 3C), and within the range of affinities previously observed in γδ TCR interactions with MHC-like^19–22^ and butyrophilin-like^2,23^ molecules. Similar results were observed between TCR95 haplotype mutants and controls (Figure S4B). In agreement with a previous report^2^, we detected no interaction between BTNL8 and any Vγ4 TCR tested (Figures S4B and S5A). Control TCRs from Vγ9Vδ2 and MR1-reactive^24^ αβ T cells showed no binding to either BTNL3 or BTNL8 (Figures S4B and S5A–B). Although BC14.1 is classically defined as a CD1b-reactive TCR, it has been reported to respond to BTNL3/8^18^. Consistent with the sufficiency of the HV4 region for BTNL3 engagement^2,3^, we detected binding of BC14.1 to BTNL3, albeit with lower affinity than other Vγ4 TCRs (Figure 3C). Together, these data demonstrate that BTNL3 engagement by Vγ4 TCRs is necessary but not sufficient for activation, and that ligand binding affinity, as measured by SPR, is uncoupled from functional responsiveness to BTNL3/8.

To directly probe the disconnect between BTNL3 binding and the absence of detectable activation, as assessed by TCR downregulation, Nur77 expression, and NFAT activation, we examined successive layers of TCR signaling. We first measured ERK phosphorylation (pERK), a downstream readout with a lower activation threshold than the cellular responses assessed above^25,26^, to determine whether BTNL3 engagement could elicit subthreshold signaling, as described for partial agonists. We also assessed proximal hallmarks of productive TCR signaling, including TCR clustering at the immune synapse and phosphorylation of CD3ζ.

Despite a large difference in reactivity of TCR81-versus TCR95-expressing cells, co-culture of Jurkat cells expressing either TCR with HEK-BTNL3+8 cells induced significantly higher pERK than did a BTNL3/8-nonreactive Vγ8 TCR control (Figures 3D, S5C, and S5D). Haplotype mutant TCRs also induced pERK, albeit at lower levels than TCR95 and TCR81 (Figures S5D and S5E), demonstrating the specificity of the TCR–BTNL3/8 interaction and establishing that SPR-detectable binding events, which may not induce full cellular reactivity, do still translate into downstream signaling reflected by ERK phosphorylation.

In contrast, proximal signaling events were more stringently restricted. Specifically, the strongly BTNL3/8-reactive TCR95 elicited more robust synapse formation and CD3ζ phosphorylation than minimally reactive TCRs, including TCR81 and TCR95 mutants (Figures 3E–F and S5F–H).

Together, these data resolve the apparent paradox between ligand binding and cellular activation by revealing a hierarchical signaling response: BTNL3 engagement supports ERK phosphorylation, but only a subset of Vγ4 TCRs transduce sufficient proximal signals to drive full cellular activation.

### Persistent physiological BTNL3/8 engagement promotes NK-like adaptation in Vγ4 IELs

To gain insights into transcriptional programs linked to cellular reactivity to BTNL3/8, we next focused on analyzing the paired scRNA- and γδTCR-seq data from γδ IELs in relation to the cloning experiment results (Figures 1C–E and S1G–H, Methods). Consistent with other studies^14,27,28^,the number of γδIELs we collected per individual was limited and highly variable (Table S1). To strengthen the robustness of our analysis and leverage known transcriptomic variation in tissue-resident lymphocytes, we combined our data with a single-cell reference atlas on a published scANVI integration model pretrained on an atlas subset containing T cells from the gastrointestinal (GI) tract^29^ (Methods). As expected, our cells aligned well with atlas γδ T cells, allowing us to then focus on a pooled subset of 8,859 gut-resident γδ T cells from 68 individuals across 14 publications, including ours (Table S4).

These cells segregated into two relatively discrete clusters recapitulating reference atlas clusters designated as “naïve γδ T” and “γδ T” cells, the latter containing most (992/1,013) of our cells (Figure S6A–B). To assess patterns of continuous transcriptional variation not captured well by clustering, we applied topic modeling—a probabilistic soft clustering approach based on non-negative matrix factorization that can be better suited to capture complex immune cell states^30–34^. A decomposition of the scANVI-corrected count matrix using three topics revealed robust, biologically coherent gene programs reproducibly observed across individuals, including our cohort (Figures 4A and S6C–D, Methods). For each topic, we tested all expressed genes for topic-specific differential expression (DE), while controlling for donor effects (Figures 4B–C and S6E–G; Table S5; Methods), and then performed pathway enrichment analysis using rank-based gene set enrichment (Figure 4D; Table S6).

**Figure 4.**
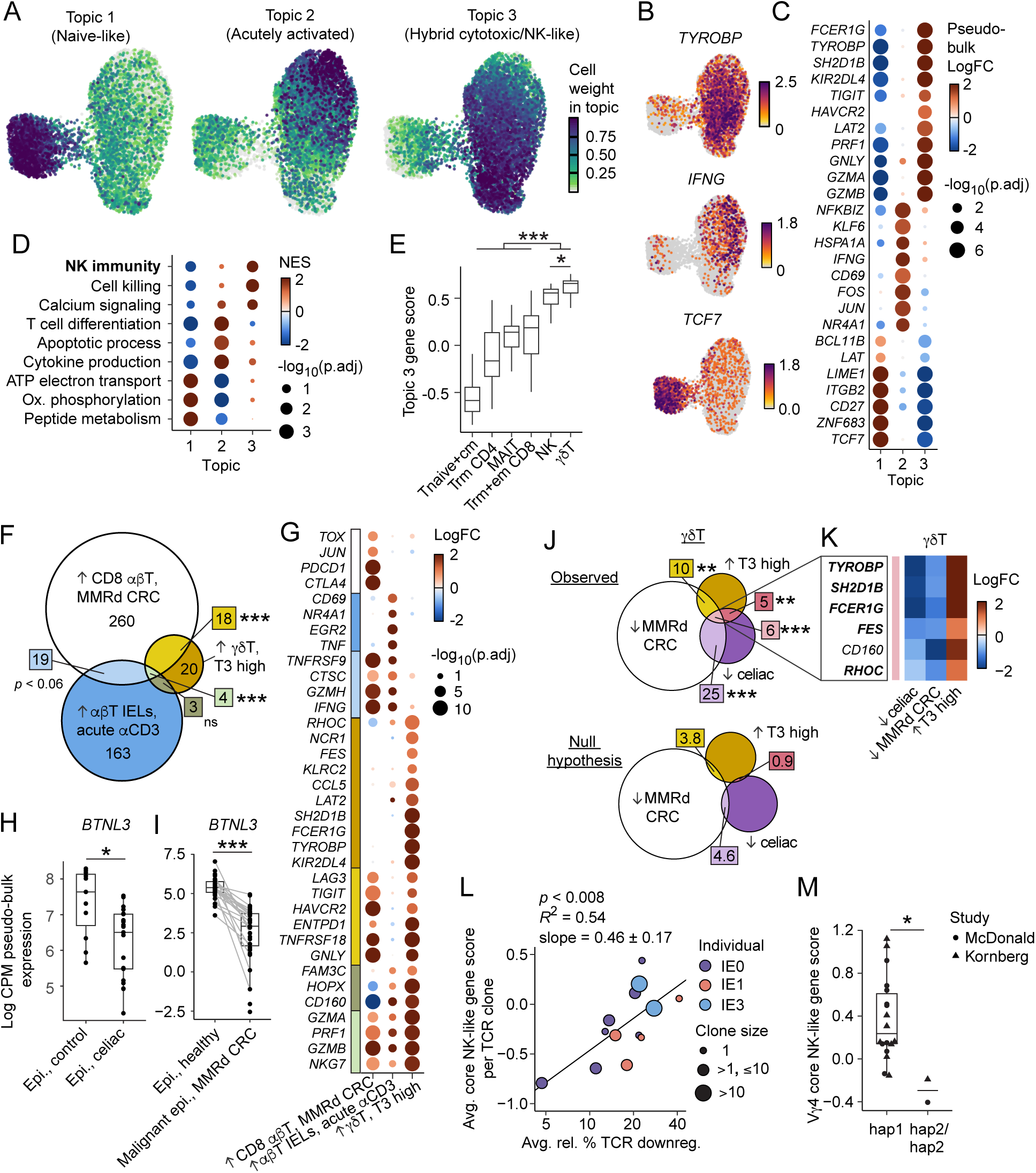
Persistent physiological BTNL3/8 engagement promotes NK-like adaptation in Vγ4 IELs. (A) UMAP embedding of gut-resident γδ T cell scRNA-seq profiles from this study and from a gastrointestinal reference atlas^29^ (*n*=8,859 cells from 68 individuals across 14 publications, including ours), colored by cell weights for three unsupervised topics identified using topic modeling^45^ on scANVI-corrected counts. (B) UMAP embedding (as in A), colored by normalized expression of a representative topic-specific DE gene for each topic (*TCF7,* Topic 1; *IFNG,* Topic 2; *TYROBP,* Topic 3). Topic-specific DE was determined by FDR-adjusted *p*<0.05 and log-fold change (logFC)>0.75, and, if gene was used in the scANVI model, by fastTopics single-cell DE with local false sign rate < 0.05 and posterior log-fold change estimate > 0.5 (Methods). (C) For curated genes (rows) DE between topics, dot plot shows relative expression (logFC, dot color) and adjusted p-values (dot size, log10 scale). Pseudo-bulk DE (determined as in B): FDR-adjusted *p*<0.05 and logFC>0.75 (Methods). (D) For curated Gene Ontology Biological Process (GO BP) pathways (rows, names abbreviated) enriched in each topic (columns), dot plot shows the normalized enrichment score (NES, dot color) and FDR-adjusted p-value (dot size, log10 scale) from rank-based gene set enrichment analysis (GSEA) (Methods, Table S6). (E) Box plots show module scores of Topic 3 genes (y axis; DE in topic 3 with FDR-adjusted *p*<0.05 and logFC>1) for curated T and NK cell subsets (x axis), computed in pseudo-bulk per individual using gene set variation analysis (GSVA) on GI reference atlas data^29^ corrected for individual-specific effects (Methods). \**p*<0.05, \*\*\**p*<0.001, Holm-corrected p-values from two-tailed Student’s *t*-test. (F) Venn diagram shows the overlaps in DE genes from three independent datasets: (1) genes higher in scRNA-seq data^37^ of CD8 T cells from biopsies of mismatch-repair deficient (MMRd) colorectal cancer, compared to that of adjacent healthy tissue (↑ CD8 αβT, MMRd CRC, white; pseudo-bulk DE: adjusted *p*<0.05; logFC>0.5); and (2) genes higher in our bulk RNA-seq data of ex vivo αβ IELs stimulated with anti-CD3, relative to that of unstimulated ex vivo αβ IELs,(↑ αβT IELs, acute αCD3; blue; DE: FDR-adjusted *p*<0.05, logFC>0.5); (3) genes higher in Topic 3 (determined as in E) (↑ γδT, T3 high; goldenrod). Shape areas are approximately proportional to the numbers of DE genes (numeric annotations) in each subset (color), including genes not expressed in all three datasets (e.g., *FCER1G*). \*\*\**p*<0.001, ns: p≥0.1; p-values from comparing observed overlap sizes to empirical null distribution, simulated from 10,000 samples (Methods; Figure S6K). (G) For curated genes (rows) DE in at least one comparison (columns, as in F), dot plot shows relative expression (logFC, dot color) and FDR-adjusted p-value (negative log scale, dot size). Annotation bar (left) colored by DE gene subset (as in F). (H) –(I) Boxplots summarize the log-normalized (log counts per million (CPM), y axis) pseudo-bulk expression of *BTNL3* in epithelial cells from public scRNA-seq datadata of: (H) the duodenum of individuals with celiac disease and of healthy controls^38^ (x axis, H), and (I) colonic regions with mismatch-repair deficient (MMRd) colorectal cancer (CRC) and adjacent healthy tissue^37^ (x axis, I). Points represent individuals; gray lines (in I) connect paired samples from the same individual. \**p*<0.05, \*\*\**p*<0.001, from limma-voom (Methods). (J) *Top*: Venn diagram shows the overlaps in DE genes from annotated γδ T cells in three independent datasets: (1) genes lower in scRNA-seq data^37^ of γδ T cells from biopsies of mismatch-repair deficient (MMRd) colorectal cancer, compared to that of adjacent healthy tissue (↓ MMRd CRC, white; pseudo-bulk DE: FDR-adjusted *p*<0.1 logFC <-0.5); (2) genes lower in scRNA-seq dat*a*^38^ of γδ T cells from the duodenal biopsies of individuals with active celiac disease, compared to that of healthy controls (↓ celiac; purple; pseudo-bulk DE: FDR-adjusted *p*<0.1, logFC < -0.5); and (3) genes higher specifically in Topic 3 (determined as in E) (↑ γδT, T3 high; goldenrod). *Bottom*: Venn diagram shows estimated expected overlaps of DE genes from the three datasets, under the null hypothesis that the three DE gene sets are drawn from independent binomial distributions (Methods). Shape areas are approximately proportional to the numbers of DE genes (numeric annotations) in each subset (color). \*\**p*<0.01, \*\*\**p*<0.001; observed overlap sizes were compared to empirical null distribution, simulated from 10,000 samples (Methods). (K) Heatmap shows relative expression (logFC, color) for genes (rows) DE in all three comparisons (columns, as in J). Genes with bolded text are also unique to the “T3 high” category in F–G. (L) Scatter plot shows the BTNL3/8 reactivity (clonally averaged relative % TCR downregulation, x axis, log-scaled to reduce the influence of outlying data) versus corrected “core NK-like” module score (y axis, Methods) of Vγ4 IEL-derived TCRs (points; point size: “clone size”, i.e., number of cells with that TCR) from 3 healthy individuals (colors). Reactivity is averaged across two experiments, with 25,000 and 100,000 HEK-BTNL3+8 stimulator cells, respectively. Module scores are corrected for an observed effect due to clonal expansion (y axis; Figure S7C; Methods). *p*<0.008 from an LRT comparing a linear model accounting for TCR downregulation and clonal expansion (black line, coefficient of determination R^2^) to a reduced model accounting only for clonal expansion (Figure S7C; Methods). (M) Boxplots summarizing the pseudo-bulk corrected “core NK-like” module scores (y axis, calculated using GSVA; Methods) of Vγ4 cells from Vγ4 hap2/hap2 individuals (x axis, right) versus from those with at least one copy of Vγ4 hap1 (x axis, left), using individuals from this study (circle) and Kornberg *et al*^14^ (triangle). \**p*<0.05, from an LRT comparing a mixed linear model accounting for haplotype and publication to a reduced model only considering publication (Methods). Statistical analysis described per panel. Nested black bars indicate statistical significance for each pairwise comparison. For boxplots (shown if *n*>3), lower/middle/upper limits of boxes: first/second/third quartiles; whiskers extend to most outlying data point or 1.5x interquartile range, whichever is closest to median. For *n*≤3, horizontal bar: median. See also Figures S6–7 and Tables S1 and S4–6.

Consistent with its strong overlap with the “naive γδ T” cluster, Topic 1 was characterized by developmental and stem-like features, including expression of the transcription factors *TCF7* (TCF-1), *ZNF683* (HOBIT), and *BCL11B*, the integrin *ITGB2*, and enrichment of metabolic pathways associated with quiescent lymphocytes (Figures 4A–D). In contrast, Topic 2 displayed features of acute TCR activation, including immediate early response genes characteristic of acute TCR stimulation (*NR4A1* (Nur77), *JUN*, *FOS* and *CD69*), together with enrichment of pathways related to T cell differentiation and cytokine production (Figures 4A–D). Distinct from both Topics 1 and 2, Topic 3 was defined by a hybrid program combining classical cytotoxic effector molecules (*GZMA*, *GZMB*, *PRF1*, *GNLY*) with NK-lineage-associated adaptor, signaling, and receptor molecules (*FCER1G*, *TYROBP*, *SH2D1B* (EAT-2), *KIR2DL4*), as well as enrichment of natural killer cell-mediated immunity pathways (Figures 4A–D). Based on these associations, we refer to the topics as “Topic 1 (naïve-like)”, “Topic 2 (acutely activated)”, and “Topic 3 (hybrid cytotoxic/NK-like)”, respectively.

Because Topic 3 (hybrid cytotoxic/NK-like) contained both cytotoxic- and NK-associated genes, we next sought to understand its expression in the context of other gut-resident lymphocyte populations. We first compared pseudo-bulk scores of the set of genes highly DE in each topic (pseudo-bulk adjusted *p* < 0.05, log-fold change > 1; “topic genes”) across gut-resident lymphocyte populations from the GI reference atlas^29^ (Methods). We found that γδ T cells exhibited the highest Topic 3 (hybrid cytotoxic/NK-like) gene scores, followed closely by NK cells, which was not the case for Topics 1 (naïve-like) or 2 (acutely activated) (Figure 4E and S6H). Notably, γδ T cells and NK cells uniquely shared expression of NK-associated adaptor molecules (*FCER1G*, *TYROBP*, *SH2D1B*, *LAT2*) and NK receptors (*KIR2DL4*, *NCR1*) (Figures 4E and S6I). Although expression of certain Topic 3 (hybrid cytotoxic/NK-like) genes (e.g., *GZMA, GZMB*) was detected in αβ T cell subsets and MAIT cells, these cell types lacked expression of the full program. For example, conventional cytotoxic tissue-resident T cells did not express key ITAM-bearing adaptor molecules such as *FCER1G* and *TYROBP* (Figure S6I), which enable cell surface expression and signaling of receptors such as CD94/NKG2C capable of activating SYK/ZAP70-dependent pathways independently of the TCR, thereby relaxing TCR-imposed specificity over key effector functions^35,36^. These findings indicate that Topic 3 does not simply reflect a generic cytotoxic state but additionally encompasses a specialized NK-like effector program associated with ITAM-coupled signaling machinery.

We next asked whether the hybrid cytotoxic/NK-like Topic 3 was associated with BTNL3/8 responsiveness. Scoring individual γδ IELs based on the expression of Topic 3 (hybrid cytotoxic/NK-like) genes, then averaging scores across cells sharing the same TCR, revealed a strong linear relationship with experimentally measured BTNL3/8 reactivity, whereas genes from Topics 1 (naïve-like) and 2 (acutely activated) showed no such association (Figure S6J, Methods). These findings established, in an unsupervised and unbiased manner, a direct connection between chronic BTNL3/8 responsiveness and acquisition of the hybrid cytotoxic/NK-like transcriptional state.

To further position Topic 3 (hybrid cytotoxic/NK-like) relative to transcriptional programs induced by acute versus chronic T cell stimulation, we compared Topic 3 (hybrid cytotoxic/NK-like) genes with signatures derived from two independent contexts. To model acute TCR stimulation, we generated bulk RNA-seq data from healthy ex vivo CD8 αβ IELs with or without acute anti-CD3 stimulation, and identified genes specifically upregulated in the acute stimulation condition (“acute TCR stimulation genes”; Methods). To characterize chronic antigen exposure in cancer, where it is associated with exhaustion, we re-analyzed a public scRNA-seq data^37^ that profiled mismatch-repair–deficient colorectal cancer (MMRd CRC) samples, along with healthy tissue controls. Using the GI-atlas pretrained model, we annotated cell types in this dataset and identified genes enriched in CD8 T cells in MMRd CRC, relative to CD8 T cells in healthy tissue (“CD8 T MMRd CRC genes”; Methods). Topic 3 (hybrid cytotoxic/NK-like) significantly overlapped with the intersection of acute TCR stimulation genes and CD8 T MMRd CRC genes, consistent with shared cytotoxic features such as *GZMA* and *GZMB* (Figures 4F, 4G, and S6K). Independently of acute TCR stimulation, Topic 3 (hybrid cytotoxic/NK-like) selectively overlapped with chronic tumor-associated programs found in the CD8 T MMRd CRC gene set, including co-inhibitory receptors such as *TIGIT*, *HAVCR2* (TIM-3), and *ENTPD1* (CD39; Figures 4F, 4G, and S6K). However, Topic 3 (hybrid cytotoxic/NK-like) notably lacked canonical exhaustion markers, such as *PDCD1* (PD-1) and *TOX,* present in CD8 T MMRd CRC genes. Instead, it was uniquely enriched for NK-associated adaptor and receptor genes (*FCER1G*, *SH2D1B*, *LAT2*, *KIR2DL4*), suggesting that chronic physiologic ligand exposure drives NK-like functional adaptation rather than classical T cell exhaustion.

Given the strong association between BTNL3/8 responsiveness and Topic 3 (hybrid cytotoxic/NK-like), we next hypothesized that disease settings characterized by reduced epithelial BTNL3/8 expression would also exhibit loss of Topic 3 (hybrid cytotoxic/NK-like) gene expression. Through re-analysis of public scRNA-seq datasets from MMRd CRC^37^ and celiac disease^38^, we confirmed reduced epithelial expression of *BTNL3* and *BTNL8* in those contexts (Figures 4H–I and S7A–B). Next, again using the GI-atlas model, we annotated cell types and isolated the set of genes with significantly reduced expression in γδ T cells in each disease context relative to γδ T cells in healthy tissues from the same dataset. Six Topic 3 (hybrid cytotoxic/NK-like) genes were downregulated in γδ T cells across both diseases, representing a significant overlap among these three gene sets (Figures 4J–K). Further, five of the six genes, *TYROBP*, *FCER1G*, *SH2D1B*, *FES*, and *RHOC*, were shared with the genes unique to Topic 3 (hybrid cytotoxic/NK-like), compared to acute TCR stimulation and exhaustion (Figures 4G and 4K; shared genes bolded in Figure 4K). Strikingly, these genes were almost exclusively NK-associated components, including the adaptor molecules *TYROBP*, *FCER1G*, and *SH2D1B*, further supporting a direct link between chronic BTNL3/8 engagement and the acquisition of an NK-like program in Vγ4 IELs.

We asked whether these five genes, which were notably not manually curated, defined a core NK-like program more tightly coupled to BTNL3/8 responsiveness than the broader 63-gene Topic 3 (hybrid cytotoxic/NK-like) signature. Despite its highly reduced size relative to Topic 3 (*n*=5 versus *n*=63), this module (“core NK-like”) exhibited a strong linear relationship with experimentally measured BTNL3/8 reactivity in γδ IELs (Figures 4L and S7C), with a steeper slope than the full Topic 3 (hybrid cytotoxic/NK-like) program, suggesting a tighter coupling to ligand reactivity. This association was particularly robust within individual donors (Figures S7D–E), demonstrating that individual-level variation in BTNL3/8 reactivity and NK-like expression were not solely driving this relationship.

Finally, we tested the relationship between BTNL3/8 reactivity and acquisition of the core NK-like program in an independent setting of genetically determined variation in BTNL3/8 responsiveness^13^. Limited sample size notwithstanding, Vγ4 IELs from *TRGV4* hap2/hap2 individuals, which encode TCRs with reduced responsiveness to BTNL3/8^13^, exhibited decreased expression of a pseudo-bulk core NK-like module score compared to hap1 carriers (Figure 4M), consistent with BTNL3/8 reactivity driving acquisition of the core NK-like gene program. In contrast, broader topic-level programs were unaffected by haplotype usage (Figure S7F). To confirm the relevance of the link between the specific core NK-like module and BTNL3/8 reactivity, we evaluated the performance of 100 random sets of 5 genes from Topic 3 (hybrid cytotoxic/NK-like) in predicting (1) experimentally measured BTNL3/8 reactivity and (2) *TRGV4* genotype. Strikingly, when considering both tasks in combination, all random gene sets performed worse than the core NK-like module (Figure S7G and Table S7), demonstrating that it is a particularly good predictor of BTNL3/8 reactivity.

Altogether, these results demonstrate that chronic physiological sensing of BTNL3/8 drives a durable NK-like program in Vγ4 IELs rather than the dysfunction or exhaustion classically associated with persistent TCR stimulation. This program is tightly linked to BTNL3/8 responsiveness across disease states and *TRGV4* haplotypes, supportive of a model in which both germline-encoded and non-germline TCR determinants couple physiological epithelial ligand engagement to chronic NK-like functional specialization in mature tissue-resident T cells.

## DISCUSSION

By integrating molecular binding, functional reactivity, genetic variation, and single-cell multiomic analyses, this study uncovers an unanticipated framework for how human intestinal Vγ4 γδ T cells respond to the epithelial butyrophilin-like heteromer BTNL3/8. Our findings revise the prevailing view that BTNL recognition is principally dictated by germline-encoded HV4-mediated binding^2–4^ and instead show that productive responsiveness is shaped by tissue selection, additional TCR determinants, and a signaling checkpoint that uncouples ligand engagement from full activation. More broadly, we show that chronic physiological ligand sensing can drive durable NK-like functional specialization rather than the dysfunction or exhaustion classically associated with persistent TCR stimulation of conventional T cells. We identify a specialized NK-like transcriptional state tightly linked to productive BTNL3/8 reactivity and marked by the expression of core adaptor molecules that license activating NK receptor signaling. This mode of peripheral functional instruction occurs after tissue entry and is therefore distinct from the thymic acquisition of effector programs by innate-like T cell subsets such as NKT and MAIT cells during development. These results provide new insights into the distinct ways in which, relative to other tissue-resident lymphocytes, γδ T cells reconcile long-term residence at barrier surfaces with the maintenance of effector capacity.

A key instigating observation was that not all Vγ4 γδ T cells are functionally reactive to BTNL3/8^7^. By cloning Vγ4 TCRs from tissues lacking BTNL3/8 expression, including thymus and blood, we found that BTNL3/8-reactive receptors were rare outside the intestine, indicating that reactivity is not an intrinsic property of all Vγ4 TCRs but instead requires selection within tissues expressing the ligand. Thus, the intestinal epithelium does not appear to simply recruit all preconfigured Vγ4 cells, but rather to shape a compartment enriched for Vγ4 γδ T cells capable of responding to BTNL3/8.

At the molecular level, our data are consistent with an important role for the HV4 region in BTNL3/8 binding and reactivity, while revealing a substantially more complex regulatory landscape governing Vγ4 responsiveness. We found that, in addition to germline-encoded polymorphisms within and beyond HV4, the non-germline-encoded CDR3γ contributed to Vγ4 responsiveness. Moreover, we identified an unexpected role for the TCRδ chain, particularly the CDR3δ. Together, these findings are consistent with a recently published Vγ4⁺ TCR–BTNL3/8 structure^39^, in which germline-encoded Vγ4 elements are the major determinant of BTNL3 recognition, whereas the associated TCR dimerization interface is centered on the CDR3γ and CDR3δ loops. Sequence or structural variation within CDR3γ, CDR3δ, or other non-HV4 regions is therefore likely to alter this dimerization interface in ways that influence the capacity of the γδ TCR to couple ligand engagement to CD3 signaling. Supporting this mechanistic model, our signaling data show that whereas HV4-dependent BTNL3 recognition was sufficient to support proximal ERK activation, productive downstream activation additionally required specific CDR3γ and CDR3δ configurations, suggesting that structural variation outside HV4 primarily governs signaling competence rather than ligand recognition. Testing this integrated structural-functional model through complementary structural and biophysical studies will be an important direction for future work.

A striking physiological implication is that the distinctive signaling architecture of γδ T cells may provide a physiological solution to chronic receptor engagement. Although all tested Vγ4 TCRs bound BTNL3, binding alone frequently failed to trigger full activation, indicating that individual ligand recognition events can generate a partial agonist-like state which is uncoupled from complete signaling output. This dissociation is consistent with recent studies showing that γδ TCR-ligand interactions, including in γδ TCR-transduced systems, are not always sufficient to drive robust signaling^19,20,40,41^. Previous work has suggested that this property reflects the greater conformational flexibility of the γδ TCR-CD3 complex, which may reduce ligand sensitivity and downstream signal propagation relative to αβ T cells^42^. Recently published γδ TCR-ligand structures that demonstrate dimeric TCR forms bound to ligand^39,43,44^ offer a complementary explanation, in which ligand binding and dimerization could both be required for strong CD3 signaling. Similar uncoupling between binding and downstream signaling has also been observed in other γδ TCR systems, including MR1-reactive receptors capable of diverse Vγ-Vδ pairings^19^. Our findings suggest that this implicit additional signaling checkpoint allows γδ T cells recognizing a constitutive, non-polymorphic self-ligand to retain adaptive capacity across distinct tissue contexts. This layered recognition logic may allow Vγ4 γδ T cells to dynamically adapt to environments in which BTNL3/8 expression is either maintained or lost, coupling constitutive tissue sensing to context-dependent functional specialization rather than fixed activation.

We furthermore propose that this two-step logic enables γδ T cells to sustain chronic engagement of non-polymorphic tissue ligands without progressing to the exhaustion programs characteristic of chronically stimulated αβ Trm cells in tumors. Instead, heightened BTNL3/8 reactivity drives differentiation into a distinct NK-like state marked by induction of *FCER1G*, *TYROBP* (DAP12), and *SH2D1B* (EAT-2). These NK-associated adaptor molecules are not induced by acute TCR stimulation, are absent from conventional CD8⁺ αβ T cells, and are lost in disease states with reduced BTNL3/8 expression, such as celiac disease, or in Vγ4 haplotypes with diminished ligand reactivity. Hence, they indicate a specialized program, rather than a generic cytotoxic response, that links chronic γδ TCR signaling to the activation of specific NK receptor pathways, which can enable TCR-independent induction of key effector functions^35,36^. This state parallels NK-like adaptations observed during chronic activation of αβ T cells in settings such as celiac disease^7^ and chronic CMV infection^6^, yet it remains fundamentally distinct from exhaustion.

Together, our findings uncover a previously unrecognized framework for γδ T cell tissue adaptation in which chronic sensing of a constitutive epithelial ligand is actively decoupled from exhaustion. We show that BTNL3/8 responsiveness is not simply hardwired through germline-encoded HV4-mediated recognition but instead emerges through a layered process integrating tissue-dependent selection, adaptive-like non-germline γδ TCR features, and an additional signaling checkpoint that uncouples ligand engagement from full activation. In this framework, germline-encoded recognition provides a foundation for tissue sensing, whereas adaptive receptor diversification selectively licenses productive responsiveness and acquisition of a specialized NK-like tissue state that preserves effector competence while partially relaxing classical TCR constraints. More broadly, these findings establish chronic physiological self-ligand sensing as a driver of adaptive tissue specialization rather than dysfunction, revealing a mode of barrier immune regulation fundamentally distinct from conventional αβ T cell paradigms.

## RESOURCE AVAILABILITY

## Lead contact

Requests for further information and resources should be directed to and will be fulfilled by the lead contact, Bana Jabri.

## Materials availability

This study did not generate new unique reagents.

## Data and code availability

- scRNA-seq and scTCR-seq data have been deposited at the Gene Expression Omnibus as GSE294419 and will be publicly available as of the date of publication.
- All original code has been deposited at Zenodo with DOI 10.5281/zenodo.14531358 and will be publicly available as of the date of publication.
- Any additional information required to reanalyze the data reported in this paper is available from the lead contact upon request.

## ACKNOWLEDGEMENTS

We thank members of the Jabri and Riesenfeld labs, Luis Barreiro, and Cezary Ciszewski for advice and helpful discussions, Adam Kornberg and Arnold Han for supplemental details regarding their data, Damien Pilotto for cell sorting, and the University of Chicago Functional Genomics Facility for RNA sequencing support. B.D.M was supported by NIH grant T32DK007074-47. H.D.A. was supported by NIH grant T32GM007183 and the National Science Foundation Graduate Research Fellowship Program under grant number 2140001. D.A.P. was supported by a Wellcome Trust Senior Investigator Award (100326/Z/12/Z). S.J.R. is a Biohub Investigator. This study was supported by an Australian Research Council discovery project (DP230102073) to B.S.G and NIH grants RC2DK133947 and R01DK067180 to B.J and R35GM138150 to A.K.

## AUTHOR CONTRIBUTIONS

B.D.M., T.M., and B.J. conceived the study. B.D.M., H.D.A., T.M., S.J.R., and B.J. designed experiments. B.D.M, T.M., C.L.S., M.J., and V.L. performed experiments. B.D.M., H.D.A., and T.M. analyzed and interpreted data. H.D.A. performed scRNA-seq, TCR repertoire, and statistical analysis. S.J.R. supervised scRNA-seq, TCR repertoire, and statistical analysis. C.L.S. and B.G. analyzed and interpreted SPR experiments. B.G. and J.R. supervised SPR experiments. M.E.B collected bulk RNA-seq data of CD8^+^ IELs. C.K., N.H., and A.K. contributed thymus samples. K.L. and J.E.M. performed primary sequence analysis of TCRs 95, 94, and 81. D.A.P. supervised sequence analysis of TCRs 95, 94, and 81. B.D.M. and H.D.A. drafted the manuscript. B.G., J.R., S.J.R., and B.J. critically edited the manuscript. D.A.P., J.R., S.J.R., and B.J. acquired funds to support the work. S.J.R. and B.J. directed the study. All authors reviewed and approved the final manuscript.

## DECLARATION OF INTERESTS

The authors declare no competing interests.

## SUPPLEMENTAL INFORMATION

Figures S1–S7.

Table S1. Number of cells captured in scRNA-seq and scTCR-seq experiments of healthy IELs with TCR usage information, related to Figures 1, 2, 3, and 4.

Table S2. TCR sequences used for cellular reactivity assays, related to Figures 1 and 4. Table S3. γδ TCR repertoire analysis, related to Figure 2.

Table S4. Number of filtered γδ T cells per study and individual in the scRNA-seq query-reference analysis, related to Figure 4.

Table S5. DE tests comparing gene expression in identified topics, related to Figure 4.

Table S6. Rank-based gene set enrichment analysis on DE tests comparing identified topics, related to Figure 4.

Table S7. Performance of random subsets of Topic 3 genes at predicting BTNL3/8 reactivity and *TRGV4* genotype, related to Figure 4.

**Figure S1. TCRs selected for BTNL3/8 reactivity assays are representative of the observed Vγ4 TCR repertoire in their respective tissue compartments, related to Figure 1**.

(A) –(B). For all (top panel, gray) or selected (bottom panel, red) Vγ4 TCRs from IELs in healthy tissue, the histograms show amino acid (AA) sequence lengths of CDR3δ (left) and CDR3γ (right) sequences (A), and clonal frequency (i.e., the fraction of an individual’s Vγ4 repertoire described by each clone) (B).

(C) Bar plots show the fraction of Vγ4 TCRs from healthy IELs whose sequences include the indicated Vδ (left), Jδ (middle), or Jγ (right) gene segments (x axis), among either cloned TCRs (red, bottom) or all detected Vγ4 TCRs (gray, top).

(D) Analogous to A, for all Vγ4 TCRs from the thymus. All clones from the thymus were detected in only one cell.

(E) – (F) Analogous to A and B, respectively, for Vγ4 TCRs cloned from PBMCs. In the top panel of (F), the y-axis is broken so that the small number of clones with higher clonal frequencies are visible.

(G) Representative flow plots show frequency of TCR downregulation in Jurkat lines expressing Vγ4 TCRs cloned from the thymus (“Thymus TCR1”), PBMCs (“PBMC TCR25”), healthy IELs (“Healthy IEL TCR6”), and active celiac disease IELs (“ACD IEL TCR92”), co-cultured with HEK control (top) or HEK-BTNL3+8 stimulator (bottom) cells (as in Figure 1C; black contours). TCR downregulation with plate-bound anti-CD3 stimulation is overlaid (gray contours, same on top and bottom).

(H) Representative flow plots show Nur77 expression in Jurkat lines expressing Vγ4 TCRs cloned from the thymus (“Thymus TCR3”), PBMCs (“PBMC TCR23”), healthy IELs (“Healthy IEL TCR13”), and active celiac disease IELs (“ACD IEL TCR90”), co-cultured with HEK control (top) or HEK-BTNL3+8 simulator (bottom) cells (as in Figure 1C).

**Figure S2. Haplotypic variants of *TRGV4,* consisting of variable amino acids within and outside of the HV4γ region, show reduced BTNL3/8 reactivity, related to Figure 1**.

(A) Amino acid sequences of naturally occurring (top rows; hap1, hap2, and hap3) and mutant (bottom rows; hap2^HV^^4^ ^hap1^ and hap1^HV4^ ^hap2^ ^or^ ^3^) Vγ4 haplotypes. Column shading and IMGT unique amino acid (AA) number (top row) indicate residues that are not consistent across all sequences with Vγ4 hap1. These residues appear in color if they differ from hap1, i.e., if they are unique to hap3 (blue), or are present in hap2 and possibly shared with hap3 but not with hap1 (red).

(B) Ribbon diagram depicting the structure of the γδ TCR of a Vγ4 hap1-encoding TCR (7RYL.pdb^20^) (inset) with an exploded view of residues in the complementarity determining regions (CDR) 1, 2, and 3, and HV4γ region (indicated by color). Key features of the locations of amino acids impacted by haplotypic variants (colored balls, annotated by IMGT unique AA number) include proximity to TCRδ chain (G28, G31), solvent-exposed side chains (S64, K85), and proximity to the HV region (Q45).

(C) BTNL3/8 reactivity (relative % TCR downregulation) in Jurkat cells expressing natural TCR95 (black) or Vγ4 mutant sequences (panel titles, red; as in A), quantified by comparing co-cultures with variable numbers of HEK control or HEK-BTNL3+8 stimulator cells (x axis). Jurkat cells expressing Vγ4 mutant sequences were also assessed by co-culturing with various concentrations of anti-CD3 (gray). Holm-corrected p-values from one-tailed LRTs of linear models using minimum and maximum HEK cell numbers as independent variables, with or without TCR labels (Methods).

(D) Plots show relative % Nur77 expression as an alternative measure of BTNL3/8 reactivity (y-axis; Methods; as in Figure 1D) for natural TCR95 with Vγ4 hap1 (black) and various TCR95 mutants (panel titles, red; as in A), co-cultured with 6,000 or 25,000 HEK-BTNL3+8 cells (x-axis). Holm-corrected p-values from one-tailed LRTs of linear models using HEK cell numbers as independent variables, with or without TCR labels.

(E) Boxplots show the fraction of cells using hap1 as in Figure 1F, but including the outlying blood patient (PB15). *\*p*<0.05, Wald test, beta regression model, as in Figure 1F (Methods; cell numbers per individual and haplotype in Table S1).

(F) Plot shows the fraction of Vγ4 cells (y axis, as in E) from PBMC samples (text labels) versus Simpson unbiased diversity (y-axis) calculated from the distributions of the fractions of Vγ4 cells that use each Vγ4 TCR in each sample (i.e., clonal frequencies).

For boxplots (shown if *n*>3), lower/middle/upper limits of boxes: first/second/third quartiles; whiskers extend to most outlying data point or 1.5x interquartile range, whichever is closest to median. For *n*≤3, horizontal bar: median.

**Figure S3. Specific γδ TCR traits are associated with tissue residence and reactivity to BTNL3/8, related to Figure 2**.

(A) –(C) Enrichment of CDR3γ motifs and sequence length: (A) Scatter plot shows the results (odds ratios, x axis, log_10_ scale; Holm-corrected p-values, y axis) of two-tailed Fisher’s exact tests of enrichment of CDR3γ motifs (points) in Vγ4 hap1 TCR clones from healthy IELs versus those from the thymus (Methods). Dashed line: threshold (0.05) for significance.

(B) Barplots show the frequencies of curated CDR3γ motifs (panel titles) in TCR clones from different tissues (x axis); these motifs were significantly enriched in IELs: \**p*<0.05, Holm-corrected two-tailed Fisher’s exact tests, as in A.

(C) Violin plots show the distributions of CDR3γ amino acid (AA) sequence lengths in Vγ4 hap1 TCR clones from the thymus and healthy IELs (x axis). ns: not significant (*p*≥0.05), two-tailed Student’s *t*-test of the log of sequence lengths.

(D) –(F) Enrichment of CDR3δ motifs and sequence length: (D) Scatter plot (analogous to A) shows the results of testing CDR3δ motifs for enrichment in Vγ4 hap1 TCR clones from the thymus versus those from healthy IELs.

(E) Barplot (analogous to B) shows frequencies of curated, IEL-enriched CDR3δ motifs (panel titles) in TCR clones from different tissues (x axis). \**p*<0.05, \*\**p*<0.01, Holm-corrected two-tailed Fisher’s exact tests, as in D.

(F) Violin plots (analogous to C) show the distributions of CDR3δ AA sequence lengths. *\*p*<0.05, two-tailed Student’s *t*-test of the log of sequence lengths.

(G) Left, the natural (“TCR94”) and CDR3δ mutant (“TCR94mut”) CDR3 AA sequences are displayed, with glycine insertions in the CDR3δ highlighted (goldenrod). Right, boxplots show BTNL reactivity (relative % TCR downregulation) of Jurkat lines expressing either TCR94 (black) or TCR94mut (goldenrod), as determined by co-culturing with variable numbers (bottom x axis) of either HEK control or HEK-BTNL3+8 stimulator cells. The relative % TCR downregulation of TCR94mut stimulated with various concentrations of anti-CD3 (top x axis) is also shown (gray). \*\*\**p*<0.001, one-tailed LRT of linear models using minimum and maximum HEK cell numbers as independent variables, with or without TCR labels (Methods).

Statistical analysis described per panel. For violin plots (in C and F), diamond: median; horizontal black bands: 25th and 75th percentiles. For boxplots (in G, shown if *n*>3), lower/middle/upper limits of boxes: first/second/third quartiles; whiskers extend to most outlying data point or 1.5x interquartile range, whichever is closest to median. For *n*≤3, horizontal bar: median.

**Figure S4. Haplotypic variants of *TRGV4* show comparable binding to BTNL3, related to Figure 3**.

(A) Plots shows % TCR downregulation (% TCR^low^, y-axis) for TCR95, TCR94, and TCR81, stimulated with various concentrations of anti-CD3 (x axis; Methods).

(B) SPR sensorgrams and derived equilibrium binding affinity curves for natural TCR95, TCR95 mutant sequences, CD1b-reactive Vγ4 TCR BC14.1, and MR1-reactive αβTCR AF-7 (columns). Each TCR was serially diluted and tested for binding with immobilized BTNL3, BTNL8, and CD1b (rows) at a surface density of ∼2000 response units. Representative sensorgram from three independent experiments each conducted in replicate (*n*=2) were used to determine the dissociation constants (K_D_) from a one-site specific binding model. Equilibrium binding curves show mean binding for replicates or no binding (NB) where undetected.

**Figure S5. Haplotypic variants of *TRGV4* show partial ERK induction but no CD3 clustering or CD3z phosphorylation in response to BTNL3/8, related to Figure 3**.

(A) SPR-derived equilibrium binding affinity curves for naturally occurring γδ TCRs (TCR95, TCR94, and TCR81), a Vγ4 CD1b-reactive TCR (BC14.1), and a control Vγ9Vδ2 TCR (as in Figure 3C). Each TCR was serially diluted from 100–0 µM and tested for binding with immobilized BTNL8 at a surface density of ∼2000 response units. Equilibrium binding curves show no binding (NB) where undetected.

(B) SPR sensorgram (left) and derived equilibrium binding affinity curves (right) for the control MR1-reactive αβTCR AF-7, performed in the same assay as Figure 3C. The TCR was serially diluted and tested for binding with immobilized BTNL3, BTNL8, and CD1b at a surface density of ∼2000 response units. Representative binding curves show no binding (NB) where undetected.

(C) Flow plots (representative of 6 independent experiments) display frequencies of TCR downregulation (as in Figure 1A) of a Jurkat line expressing IEL-derived Vγ8 TCR40 (as in Figure 3D) in co-cultures with 10^5^ HEK control (top) and HEK-BTNL3+8 stimulator (bottom) cells (black contours). TCR downregulation with plate bound anti-CD3 stimulation is overlaid (gray).

(D) Representative flow plots show pERK expression (y axis) in Jurkat lines expressing Vγ4 TCRs (TCR95 hap1, TCR81TCR81, TCR95 hap2, TCR95 hap3) or a Vγ8 TCR (TCR40) (columns), co-cultured (as in Figure 1C) with HEK control (top) or HEK-BTNL3+8 simulator (bottom) cells.

(E) Plots show relative % pERK^hi^ (y axis, Methods) for Jurkat lines expressing BTNL3/8-reactive natural Vγ4γ hap1 TCR95 (orange), haplotype mutant sequences (blue, purple) or a BTNL3/8-non-reactive Vγ8 TCR (TCR40, gray), each cultured as in Figure 1C. Horizontal bars represent median. For each parameter combination, *n*=4–6. \*\**p*<0.01, \*\*\**p*<0.001, Holm-corrected p-values from one-tailed LRTs of linear models using minimum and maximum HEK cell numbers as independent variables, with or without TCR labels (Methods).

(F) –(H) Representative images (F) and summary quantification in violin plots of CD3 (G) and pCD3ζ (H) staining of Jurkat lines expressing TCR95 hap1, TCR95 hap1→hap2, and TCR95 hap1→hap3, co-cultured with either HEK control or HEK-BTNL3+8 stimulator cells (rows, F; x axis, G, H). \*\*\**p*<0.001, ns: not significant (*p*≥0.05); Holm-corrected p-values from a two-tailed Student’s *t* test.

Statistical analysis described per panel. Nested black brackets indicate statistical significance for each pairwise comparison. For violin plots (in G and H), diamond: median; horizontal black bands: 25th and 75th percentiles.

**Figure S6. Topic modeling identifies a gene program conserved between individuals and linked to BTNL3/8 reactivity, related to Figure 4**.

(A) UMAP embedding of gut-resident γδ T cells (as in Figure 4A), with cells colored by annotations presented in the reference publication^29^.

(B) UMAP embedding (as in Figure 4A), with γδ IELs sequenced for this study colored in red, and those from other studies colored in light gray.

(C) UMAP embeddings (as in Figure 4A), with cells sequenced for this study colored by topic weight for each topic (panel title), and cells from other studies colored in light gray.

(D) UMAP embeddings (as in Figure 4A), with cells highlighted (in red) for each individual (panel title) profiled for this study, compared to cells from other individuals (light gray).

(E) Topic weight distributions for cells from the entire query-reference dataset (light gray) and from only this study (red). Vertical dashed lines indicate thresholds for pseudo-bulk comparisons, above which cells are designated as “high” in a topic, and below which as “low” in a topic. For Topic 1 (naïve-like), the cell weight threshold was 0.3; for Topic 2, 0.125; and for Topic 3, 0.2.

(F) UMAP embeddings (as in Figure 4A), colored by normalized expression of two additional, representative, topic-specific DE genes (determined as in Figure 4B) (panels) for each topic.

(G) For each topic (T1, T2, T3), a pair of heatmaps show z-scored pseudo–bulk normalized expression (color), corrected for individual-specific effects, of selected topic-specific DE genes (determined as in Figure 4B) (rows) in pseudo-bulk samples created from cells with low (left) or high (right) topic weights (as in E, Methods); pseudo-bulk is per individual (column) from various studies (top color bar).

(H) Violin plots show, for each topic (panel title), pseudo-bulk topic module scores (y axis), calculated per individual (as in Figure 4E) from genes highly DE in each topic (as in H), after correcting for individual-specific effects in curated T and NK cell subsets (x axis) from a GI reference atlas^29^. Diamond: mean; horizontal black band: median. \**p*<0.05, \*\**p*<0.01, \*\*\**p*<0.001, Holm-corrected p-values from two-tailed Student’s *t*-test. Nested black bars indicate statistical significance for each pairwise comparison.

(I) Heatmap shows row-z-scored median expression (color) in pseudo-bulk samples, calculated after correcting for individual-specific effects, of curated Topic 3 (hybrid cytotoxic/NK-like)–specific DE genes (rows) in curated T and NK cell subsets (columns) from the GI reference atlas ^29^.

(J) For each topic (panel title), scatter plot (analogous to Figure 4L), shows the BTNL3/8 reactivity (clonally averaged relative % TCR downregulation, x axis, log-scaled to reduce the influence of outlying data) of Vγ4 IEL-derived TCRs (points; point size: number of cells in clone, i.e., with that TCR) from 3 healthy individuals (colors), versus clonally averaged topic module scores (y axis), based on genes highly DE in each topic (FDR-adjusted *p*<0.05; logFC>1) and corrected for an observed effect due to clonal expansion (y axis; Methods). Statistical significance (annotated p-value, slope) was determined (as in Figure 4L; Methods) using LRT to compare a linear model accounting for TCR downregulation and clonal expansion (black line) to a reduced model accounting only for clonal expansion (Methods).

(K) Venn diagrams (as in Figure 4F) show the average expected overlap in DE genes across the three independent datasets from Figure 4F under the null hypothesis that the three DE gene sets are drawn from independent binomial distributions (bottom). Observed overlap in DE genes (top) is reproduced from Figure 4F for clarity. \*\*\**p*<0.001, ns: p≥0.1; p-values reproduced from and calculated as in Figure 4F.

Statistical analysis described per panel.

**Figure S7. Topic modeling identifies an NK-like gene program characteristic of chronic antigen exposure, related to Figure 4**.

(A) –(B) Boxplots (analogous to Figure 4H–I) summarize log-normalized (log counts per million (CPM)) pseudo-bulk expression of *BTNL8* in epithelial cells from public scRNA-seq datasets (as described for Figure 4H–I). Points represent individuals; gray lines (in B) connect paired samples collected from the same individual \*\**p*<0.01, \*\*\**p*<0.001, from limma-voom (Methods).

(C) Scatter plot (analogous to Figure 4L) shows the BTNL3/8 reactivity (clonally averaged relative % TCR downregulation, x axis, log-scaled to reduce the influence of outlying data) versus a module score of core NK-like genes (y-axis) (as in Figure 4L), but without correction for clonal expansion. Statistical significance and coefficient of determination R^2^ as in Figure 4L. Black lines show the full linear model fit accounting for TCR downregulation (slope of both lines) and clonal expansion (difference between top and bottom line). The intercept of top line is the sum of the linear model’s estimated intercept and the coefficient modeling clonal expansion; the intercept of bottom line equals only estimated model intercept (Methods).

(D) Scatter plot (analogous to Figure 4L) shows, for single individual IE0, the BTNL3/8 reactivity (clonally averaged relative % TCR downregulation, x axis, log-scaled to reduce the influence of outlying data) versus module score of core NK-like genes (as in Figure 4L), corrected for an observed effect due to clonal expansion (y-axis; Methods), for Vγ4 IEL-derived TCRs (points; point size: number of cells in clone) from only healthy individual IE0. Statistical significance (p-values as annotated; slope) was determined as in Figure 4L, but only using TCR clones derived from IE0.

(E) Scatter plot (analogous to Figure 4L) shows the BTNL3/8 reactivity (clonally averaged relative % TCR downregulation, x axis, log-scaled to reduce the influence of outlying data) versus a module score of Topic 3 (hybrid cytotoxic/NK-like) genes (as in Figure S6H), corrected for an observed effect due to clonal expansion (y axis, Methods) of Vγ4 IEL-derived TCRs (points; point size: number of cells in clone) from only healthy individual IE0. of. Statistical significance (p-value as annotated; slope) was determined as in Figure 4L, but only using TCR clones derived from IE0.

(F) Boxplots with gene module scores analogous to those in Figure 4M, but calculated from sets of all genes highly DE in each topic (panel title; gene sets as in Figure S6J). ns: not significant, *p*≥0.05; determined as in Figure 4M (Methods).

(G) Scatter plot summarizes the associations between 100 random 5-gene subsets of Topic 3 genes (black points) and two metrics of BTNL3/8 reactivity: *TRGV4* genotype (x axis) and experimentally measured BTNL3/8 reactivity in Vg4 hap1 IELs (y axis), as compared to the core NK-like module of five genes (orange diamond; determined as in Figures 4L–M). Associations are measured by estimated slopes from linear models (Methods; analogous to models used in Figures 4L–M). Thresholds for outperforming the core NK-like module are shown for each metric independently (orange dashed lines) and for joint performance evaluated via both metrics (shaded orange rectangle).

Statistical analysis described per panel. For boxplots (shown if *n*>3), lower/middle/upper limits of boxes: first/second/third quartiles; whiskers extend to most outlying data point or 1.5x interquartile range, whichever is closest to median. For *n*≤3, horizontal bar: median.

## METHODS

### Participants

PBMCs were isolated from healthy individuals or, in a single case, from a celiac disease individual on a gluten-free diet (PB13). IELs were isolated from healthy control individuals or patients with active celiac disease undergoing standard of care endoscopic procedures. Healthy control individuals were screened to exclude patients with a history of celiac disease, inflammatory bowel disease, active malignancy, or use of immunosuppressive medications. Samples were isolated by pinch biopsy from either the duodenum or the left colon (healthy controls) or the duodenum (active celiac patients). PBMC and IEL samples were isolated from patients between 18–65 years old. Thymus fragments were obtained from 6–24-week-old patients undergoing standard of care cardiac surgery. A single patient had trisomy 21 (TY16), but all other patients studied did not have known genetic abnormalities. Samples were obtained under IRB protocols 201854, 12623B, and 15573A from the University of Chicago and protocol 2020-203 from Advocate Aurora Health. All patients or their guardians gave written consent.

### Lymphocyte Isolation

Thymus fragments were collected in the operating room and transported to the laboratory in F12 media. Thymocytes were isolated by mechanical dissociation from the surgical specimen. PBMCs were isolated using the standard Ficoll gradient method. IELs were isolated by shaking pinch biopsies at 250 rpm at 37 °C for 30 minutes in RPMI-1640 (Gibco), supplemented with 1% dialyzed FCS, 2 mM EDTA, and 1.5 mM MgCl2. To maximize recovery, a second 30-minute shake was performed in fresh media and the supernatants were combined.

### Flow Cytometry

Following isolation, cells were resuspended in PBS, supplemented with 2% FBS. The following biotinylated or directly conjugated antibodies against CD3 NovaBlue 610-70s (UCHT1), CD3 PE-Cy7 (UCHT1), CD4 biotin (A161A1), CD4 NovaBlue 660 (SK3), CD8α BV650 (RPA-T8), CD19 biotin (HIB19), CD45 AF647 (HI30), CD45 BV711 (HI30), CD103 BUV661 (Ber-ACT8), CD235ab biotin (HIR2), Nur77 PE (12.14), pCD3ζ (3ZBR4S), pERK1/2 (6B8B69), TCRαβ biotin (IP26), TCRαβ BUV737 (IP26), TCRαβ BV421 (IP26), TCR γδ APC (B1), Vδ1 APC (REA173), Vδ1 PerCP Vio700 (REA173), Vδ2 FITC (B6), Vγ4 PE (4A11.904) were purchased from Biolegend, Exbio, Invitrogen, Miltentyi, Thermo Fisher Scientific, or BD Biosciences. Intracellular Nur77 staining was performed using the True-Nuclear transcription factor buffer set (Biolegend). pERK staining was performed using BD Phosflow Fix Buffer 1 (557870) and Perm Buffer 3 III (558050). In some experiments, cells were stained with LIVE/DEAD fixable blue dye (Thermo-Fisher Scientific) to identify dead cells. To enrich for γδ T cells prior to sorting, thymocytes were stained with biotin-conjugated antibodies against CD19, CD235ab, and CD4, and PBMCs were stained with biotin-conjugated antibodies against CD19, CD235ab, and TCRαβ. Cells were washed and then incubated with magnetically labeled streptavidin beads (Biolegend). Labeled cells were removed with an LS column (Miltenyi). All staining was done in the dark on ice for 20-30 minutes. Samples were collected on a Cytek Aurora or BD Symphony S6 and data was analyzed using FlowJo v10.8.1.

### Jurkat cell generation

TCRγ and TCRδ chain sequences obtained from scTCR-seq of thymocytes, PBMCs, and IELs were cloned into pLV-RFP and pLV-GFP lentiviral vectors (Biosettia), respectively. TCRs were stably expressed via lentiviral transduction of TCRαβ negative Jurkat76 cells (gift of Mark Davis)^46^. Jurkat lines were cultured in RPMI-1640 supplemented with 10% FBS, 2mM Glutamax, and penicillin/streptomycin. After 72 hours of recovery, Jurkat lines were used in BTNL3+8 reactivity assays.

### Ex vivo HEK-BTNL3+8 reactivity assay

18 hours prior to stimulation, 10^5^ HEK or HEK-BTNL3+8 cells were placed in 96-well round bottom plates as previously described^7^. Following lymphocyte isolation, cells were washed in RPMI supplemented with 10% FBS, 2mM Glutamax, and penicillin/streptomycin. 10^5^ lymphocytes were overlaid onto the HEK cell monolayers and cultured for 24 hours at 37 °C prior to analysis of CD3 (UCHT1) and Vγ4 (4A11.904) downregulation by flow cytometry.

### In vitro HEK-BTNL3+8 reactivity assay

Eighteen hours prior to stimulation, 96-well round-bottom plates were coated with 10^5^, 2.5x10^4^, or 6.25x10^3^ HEK or HEK-BTNL3+8 cells. The following morning, 5x10^4^ transduced Jurkat cells were added on top of the HEK cell monolayer and cultured at 37 °C for 2 hours prior to analysis of CD3 (UCHT1) and TCRγδ (B1) downregulation by flow cytometry. As a positive control, Jurkat lines were incubated for 2 hours with 0.5, 1, 2, or 4 ug/mL of plate bound anti-CD3. A Jurkat line expressing a highly BTNL3/8 reactive TCR (IE95) was tested during each experiment as an additional positive control. While maximal cell surface TCR expression varied between Jurkat lines, there was not a relationship between the MFI of the transduced TCR and BTNL3/8 reactivity.

For experiments that quantified Nur77 upregulation, stimulation cultures were set up as above, but only the 2.5x10^4^ and 6.25x10^3^ concentrations of HEK or HEK-BTNL3+8 cells were used. For experiments that used the NFAT-luciferase reporter, stimulation cultures were set up similarly but were incubated for 4 hours. Cells were then harvested and luciferase activity was quantified using the Nano-Glo assay (Promega). For experiments that quantified pERK induction, stimulations cultures were set up as above but were incubated at 37 °C for 30 minutes prior to analysis.

### Measuring TCR-BTNL3/8 reactivity with relative percent TCR downregulation, relative percent Nur77 upregulation, and scaled luminescence

In both the ex vivo and in vitro TCR downregulation assays, we primarily evaluated BTNL3/8 reactivity using a relative percent TCR downregulation metric that accounted for two factors: (1) the change in %TCR^low^ cells between HEK-BTNL3+8 and HEK (untransduced) conditions (representative gating strategy shown in Figure 1A), and (2) the full possible range of this change. We defined this metric by scaling (1) by (2). Specifically, for the ex vivo assay, **relative % TCR downregulation** was calculated using the following formula:

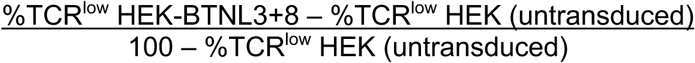

In the in vitro assay, when paired anti-CD3 and HEK measurements were available (Figures 1C, 1E, S2C, 4L, S6H, and S7C–E), we additionally took into account the positive anti-CD3 control by using it to determine the scaling for relative % TCR downregulation as follows:

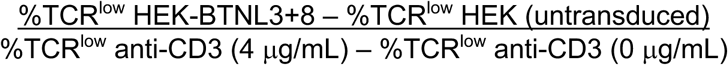

The relative % TCR downregulation upon incubation with anti-CD3 was also scaled using the full range of that experiment:

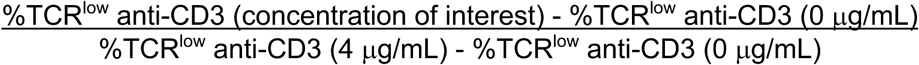

Otherwise, in Figures 2C, 2F, and S3G, for which paired anti-CD3 and HEK measurements were not available, relative % TCR downregulation was defined as in the ex vivo assay.

Similarly, **relative % Nur77 upregulation** was defined as:

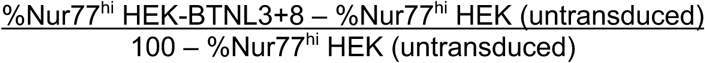

And **relative % pERKh**i was defined as:

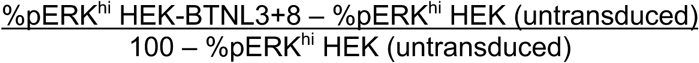

To determine **relative NFAT luminescence**, we first calculated raw relative NFAT luminescence by subtracting luminescence in untransduced HEK, as:

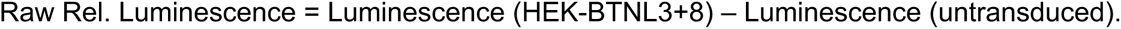

Then, these raw values were batch-corrected on the log scale. Specifically, batch coefficients were calculated by fitting the following linear model in R:

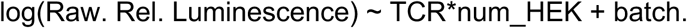

Only the maximum and minimum number of HEK cells were used for this calculation, ensuring that the data followed a linear relationship. Then, the batch terms were subtracted from all datapoints in log space (including those with 12,500 HEK cells) to present the data in Figure 3B.

### Confocal Microscopy

Coverslips were pre-treated with 0.05% poly-L-lysine (Sigma-Aldrich) for 20 minutes at room temperature, washed 3 times with sterile water, and allowed to air dry. Cultures of TCR-expressing Jurkat cells and HEK or HEK-BTNL3+8 cells were harvested, washed, and resuspended at 2 x 10^6/mL in RPMI-1640 supplemented with 10% FBS, 2mM Glutamax, and penicillin/streptomycin and pooled together in an Eppendorf tube. 8000 Jurkat cells and 4000 HEK or HEK-BTNL3+8 cells were placed on the pre-coated coverslips and incubated at 37 °C in 5% CO_2_ for 30 minutes. Coverslips were then washed with PBS, fixed with 4% paraformaldehyde for 10 minutes at room temperature, and permeabilized in PBS with 0.2% BSA and 0.05% saponin for 30 minutes at room temperature. Antibody staining was performed in permeabilization buffer for 1 hour at room temperature. Coverslips were mounted with Fluoromount-G, dried overnight, and imaged using a spinning disk confocal microscope with a 40x objective (Oxford Instruments). Image analysis was performed using QuPath (v0.5.1). Analysis was performed using a single median-plane image for each cell-cell conjugate extracted from the z-stack.

### Single-cell RNA-sequencing

For PBMCs, Vδ2+ and Vδ2-γδ T cells were sorted separately and then pooled to ensure that the more homogeneous Vδ2+ (typically Vγ9+) population accounted for approximately 30% of total cells. Because thymocyte and IEL γδ T cells were more diverse, this was not required for cells isolated from these tissue locations. Following sorting, samples from multiple individuals sorted on the same day were pooled to minimize reagent usage. Samples were processed using the 5’ single cell version 1.1 kit (10X Genomics) according to the manufacturer’s instructions with 2000–6000 cells captured from each sample. Libraries for the thymus, PBMCs, and first three IEL batches were sequenced using an Illumina Novaseq 6000 instrument, while the last batch was sequenced using an Illumina Novaseq X. The first IEL batch was frozen and spiked with 15–50% αβIELs per individual to increase cell number and minimize cell loss during processing, while the following batches were fully processed on the day of collection and not spiked with αβIELs.

### Single-cell TCR-sequencing

Following cDNA amplification, TCR libraries were generated using the 5’ single cell version 1.1 kit (10X Genomics), but with a custom set of primers, as previously described^47^. The first PCR amplification used 2 μM of forward primer 5’-AATGATACGGCGACCACCGAGATCTACACTCTTTCCCTACACGACGCTC-3’ and 1 μM each of reverse primers 5’-CCCACTGGGAGAGATGACAA-3’ and 5’-ATCCCAGAATCGTGTTGCTC-3’. The second amplification step used 2 μM of forward primer 5’-AATGATACGGCGACCACCGAGATCT-3’ and 1 μM each of reverse primers 5’-GACAAAAACGGATGGTTTGG-3’ and 5’-GGGGAAACATCTGCATCAAG-3’. Libraries were sequenced using Illumina Novaseq instruments as in the paired scRNA-seq.

### ScRNA-seq data processing and quality control

Count matrices were obtained using the *count* command from the CellRanger software (version 6.1.2)^48^. Next, the cells from multiple individuals were demultiplexed from the nucleotide polymorphisms present in their transcript reads using the Souporcell pipeline (ver. 2)^49^. Quality control filtering was then performed on an individual-specific basis using the following criteria: Souporcell-identified doublets and unassigned cells were discarded. Then, the joint distribution of the number of unique genes per cell and the number of unique transcripts per cell was visualized per individual, and outliers were discarded. Based on a qualitative examination of the plots, thresholds for calculating outlying data were set at follows: cells with less than 500 or greater than 20,000 UMIs were excluded, as were cells with less than 300 or greater than 6,000 unique genes. Similarly, the distribution of the percent of mitochondrial RNA reads of a cell’s total profiled transcriptome was visualized per individual, and outliers from this distribution were also discarded. Mitochondrial percentage thresholds for outlying were also determined by visual examination, varied by individual, and ranged from 10% to 25%. Overall, these quality control measures filtered 585/2,714 cells (21.6%) from the dataset, resulting in a total of 2,129 cells for downstream analysis.

### ScTCR-seq data processing, clonotype determination, and analysis

γδTCR information was obtained using the *vdj* command from CellRanger with an altered IMGT reference^48^. Specifically, because CellRanger V(D)J processing is specialized for αβTCR detection, the *TRG* and *TRD* gene names in the IMGT reference were modified to follow the *TRA* and *TRB* naming conventions. The custom inner enrichment primers described above were also specified. Clone assignments from CellRanger, which aim to estimate cells sharing the same paired TCR while accounting for technical limitations such as the detection of single (as opposed to paired) TCR chains in some cells, were used for downstream analysis. Next, to receive more detailed TCR information, the filtered cell-specific TCR contigs detected with *vdj* were analyzed with IMGT’s High/V-QUEST tool^50^. Only TCRs from cells which passed scRNA-seq quality control were used for downstream analysis. Of the cells which passed scRNA-seq quality control, from the batches which were not spiked with αβIELs (batches 2–4), 57% (811/1,421) had detected TCRγ and TCRδ chains, while an additional 11% (162/1,421) had either a TCRγ or TCRδ chain detected, but not both.

### Generalized linear models

To compare the relative percent TCR downregulation between TCR sequences obtained from multiple tissues in multiple individuals (Figure 1C), we used a mixed linear model with the following components: a tissue coefficient representing the mean relative percent TCR downregulation per tissue across individuals; and an individual-specific intercept representing the mean relative percent TCR downregulation for that individual. That is, the model took the form of *rel_pct_TCR_downregulation ∼ tissue + (1|individual)*, following the syntax of the lme4 R package (version 1.1-36). Significance was determined using an LRT to compare this full model with a reduced model that included only the individual-specific intercepts (i.e., *rel_pct_TCR_downregulation ∼ (1|individual)*). To determine the pairwise p-values presented in Figure 1C, separate models were fit for each combination of tissue pairs (e.g., for comparing thymus and healthy IELs, *tissue* had two categorical levels: *thymus* and *healthy_IELs*).

To evaluate the effect of TCR on BTNL3/8 reactivity metrics in experiments using multiple numbers of HEK cells (Figures S2C, S2D, 2C, S3G, 3A, 3B, 3D, and S5E), we accounted for multiple HEK cell numbers in the same formula by modeling the effects of both the cloned TCR and HEK cell number on the BTNL3/8 reactivity metric, using only the minimum and maximum HEK cell numbers so that the data could be modeled linearly. That is, we used the following linear formula in R: *BTNL38_reactivity ∼ num_HEK_cells*TCR*. To estimate statistical significance, we used an LRT comparing this linear model to a reduced model only accounting for HEK cell number (*BTNL38_reactivity ∼ num_HEK_cells*). For pairwise statistical comparisons, we fit these models using only the relevant pair of TCRs, as opposed to all TCRs tested in the experiment.

To assess the association between the average relative percent TCR downregulation and average gene module scores per clone (Figures 4L, S6J, S7C–E, and S7G), we used a linear model accounting for both relative percent TCR downregulation and an observed effect due to clonal expansion. That is, we used the formula *avg_module_score ∼ rel_pct_TCR_downregulation + is_expanded*, where *is_expanded* was *TRUE* for clones consisting of more than one cell and *FALSE* otherwise. *FALSE* was used as the reference. To reduce noise and the influence of outliers, the relative percent TCR downregulation was averaged across measurements using two different numbers of HEK-BTNL3+8 stimulator cells (25,000 and 100,000), then log-transformed. To regress out the effect of expansion (Figures 4L, S6H, S7D–E, and S7G), the coefficient estimated for *is_expanded* was added to the modules scores for unexpanded clones. To assess statistical significance of the TCR downregulation term, an LRT was performed comparing models with and without the *rel_pct_TCR_downregulation* term. That is, the full model *avg_module_score ∼ rel_pct_TCR_downregulation + is_expanded* was compared to the reduced model *avg_module_score ∼ is_expanded*.

To evaluate the influence of *TRGV4* genotype on gene module scores (Figures 4M, S7F, and S7G), we accounted for the effects of both genotype and the publication in which the data were presented by using a linear model with the formula *module_score ∼ is_hap2_homozygote + publication*, where *is_hap2_homozygote* was encoded as *TRUE* for hap2/hap2 individuals and *FALSE* (the reference level) for others. Our cohort (McDonald) was used as the reference level for publication. The presented data regresses out the effect of publication, which was highly significant (*p*<0.001, LRT), by subtracting the estimated publication coefficient from the Kornberg module scores. To assess statistical significance of the *is_hap2_homozygote* term, an LRT was performed comparing models with and without it. That is, the full model *module_score ∼ is_hap2_homozygote + publication* was compared to the reduced model *module_score ∼ publication*.

Beta regression was used to model the fraction of hap1 versus hap2 or hap3 usage in different tissues (Figures 1F and S2E) due to its suitability for modeling proportional data within the open unit interval of 0 to 1. It was performed using the *betareg* package (version 3.1-4) in R^51^. In part because two data points had proportions of 1 (explained in “*TRGV4* haplotype determination” above), which is outside the bounds of the beta distribution, all data were transformed prior to testing in a manner consistent with prior publications^52^ using he transformation function *(p*(n - 1) + 0.5)/n*, where *p* is the frequency of the haplotype and *n* the total number of Vγ4 cells per individual. Significance of hap1 enrichment in the intestine was then assessed via an LRT comparing models with and without a term considering gut residency, i.e., full model: *transformed_hap1_freq ∼ is_intestinal_sample*, and reduced model: *transformed_hap1_freq* ∼ *1*.

### *TRGV4* haplotype determination

Haplotype assignments were required to be an exact match to the protein sequences presented in Figure S2A. Infrequent *TRGV4* sequences that were not an exact match for any haplotype (fewer than 1% of total *TRGV4* TCRs) were discarded prior to downstream haplotype comparisons (Figures 1F, S7F, and S7G). An individual was considered homozygous for hap2 if all *TRGV4* cells from that individual used only hap2. No hap3/hap3 homozygotes were detected in data collected for either this publication or for Kornberg *et al*^14^. For Viswanathan *et al*^15^., no hap2/hap2 or hap3/hap3 homozygotes were detected. Similarly, individuals classified as heterozygous were required to have at least one TCR from two different haplotypes (hap1 and hap2, hap1 and hap3, or hap2 and hap3). For the Kornberg *et al.* data, following their analysis, CD103^−^ cells were assumed to be from the lamina propria and excluded from downstream analysis but were used for *TRGV4* genotyping, resulting in the two individuals called as heterozygotes having 100% hap1 CD103^+^ IELs (Figure 1F). For Viswanathan *et al.*, CD103 status of cells was unknown, and the authors did not document any isolation of the epithelial fraction. Thus, the single intestinal data point from Viswanathan *et al.* likely includes some cells from the lamina propria, though previous work has shown that a median of 90% of intestinal Vγ4 T cells are CD103^+^^10^, suggesting that most of these cells likely reside in the epithelium.

Two heterozygous samples (Thy2 from Viswanathan and CONTROL-7 from Kornberg) had very low numbers of cells with a sequenced *TRGV4* gene segment (less than 5; Table S1) but were included in the analysis to represent the observed variability in *TRGV4* usage frequency. The observed gut-specific enrichment of *TRGV4* hap1 was still significant if these samples were excluded (*p*<0.05, LRT comparing beta regression models with and without a term comparing gut residency, as discussed in “Generalized linear models”).

Due to the enrichment of hap1 cells in IELs (Figure 1F), it is possible that some individuals for whom only hap1 cells were detected were falsely classified as hap1/hap1 homozygotes and were actually hap1/hap2 or hap1/hap3 heterozygotes, particularly in our cohort, where the lamina propria was not sequenced. However, we do not expect any such errors to notably change our results, since cells from hap1 homo- and heterozygotes were treated the same in the transcriptional analysis.

### γδ TCR repertoire analysis

An initial set of CDR3 motifs was constructed from the set of all sequences of up to 20 amino acids in length present in the CDR3 sequences in the data. From these, we added motifs consisting of the same sequences with one amino acid substituted by a “wildcard” (matching any amino acid, represented by an “X”). Motifs that occurred in fewer than three individuals were discarded. The remaining motifs were then hierarchically clustered based on a Jaccard similarity matrix describing the fraction of cells in which each pair of motifs co-occurred. We clustered based on a threshold of 0.9 Jaccard similarity, after considering multiple thresholds and finding that this value performed reasonably well at grouping similar motifs without losing statistical power in downstream analysis.

Next, we tested for motif cluster enrichment using binary comparisons between two categories (i.e., thymus versus IELs). For the thymus comparison, to increase the number of individuals profiled, we pooled our data with that of a recent publication that also performed single-cell γδ TCR-seq of thymocytes^16^. Due to limited, often single-digit numbers of clones per individual in the IELs, we used Fisher’s exact tests to compare motif cluster usage between categories, which does not account for individuals. However, we did enforce some degree of robustness in the signals detected across individuals by removing all motif clusters that did not co-occur in at least three individuals in one of the two categories prior to testing. Further, to ensure that our tests were powered to detect differences between categories with vastly different clone numbers (e.g., 84 clones from the IELs vs. 1,673 from the thymus for CDR3δ), motif clusters were also required to occur with a minimum frequency that would be detectable in both categories. This frequency threshold corresponded to three clones in the IELs. For the thymus vs. IEL CDR3δ comparison, this meant that a motif cluster had to be present in roughly 3.6% of clones from the thymus or the IEL compartment (i.e., 3 out of 84 clones in the IELs). We suspect that meaningful signal was also present in less frequently occurring clones, but we were not powered to detect it.

To account for significant statistical dependences between motif clusters, p-values from the Fisher’s exact tests were adjusted using the Holm method, which controls the family-wise error rate when tests can have arbitrary dependences and is slightly less conservative than the Bonferroni correction but with the same theoretical guarantees (Figures S3A and S3D).

For the 100 permutations used to compute the empirical null p-value distribution (Figures 2A and 2D), categorical labels (i.e., thymus and IEL) were permuted without replacement using the sample function in R. To ensure that the permuted tests were equivalently powered as the tests with true labels, minimum frequency thresholds were re-applied using the new categorical labels prior to running Fisher’s exact tests on the permuted data. To focus on the signal specifically in low p-values, we compared the tails of the resulting empirical null p-value distributions to the tail of the observed p-value distribution by taking the sign of the maximum difference between the two eCDFs only for p-values less than 0.05. This was designed as an empirical equivalent to a one-sided Komologorov-Smirnov test evaluating whether the tail of the actual p-value distribution was less than the permuted distributions.

### Surface plasmon resonance

Surface plasmon resonance experiments were conducted at 25 °C on a BIAcore 3000 of T200 instrument in 20 mM HEPES (pH 7.4) and 150 mM NaCl with an addition of 0.005 % (v/v) surfactant p20 (Cytiva). BTNL3, BTNL8 and CD1b were captured on a CM5 sensor chip (Cytiva) via amine coupling, whereby ∼2000 response units of each were immobilised. Serial dilutions from 100 to 0 μM of TCRs TCR81, TCR94, TCR95, TCR95 haplotype mutants, Vγ9Vδ2, BC14.1 and AF7 were injected over the immobilised proteins at a flow rate of 5 μL.min^-1^. Two repeated experiments were conducted, each with included experimental duplicates. Sensorgram plots, equilibrium binding curves and kinetics were calculated in GraphPad Prism using a one-site specific binding model.

### Integration with the pan-GI reference atlas

To integrate our data with the public reference atlas, we used the scArches pipeline^53^ on a single-cell annotation using variational inference (scANVI) model, published with the atlas manuscript, that was trained on all T cells and innate lymphocytes^29^. Our query data consisted of individual samples from tissues that were not frozen and that contained at least 50 cells passing quality control (i.e., IE3–5, IE7, IE8, and IE10, which together include samples from all three batches; metadata published in Table S1). Including frozen samples that passed the cell quality threshold (IE0–IE2, IE9) slightly reduced the quality of the resulting embedding, and increased training time, but otherwise yielded similar results.

Next, to integrate our data into the existing published model, we trained on a concatenation of the query and reference data using the scANVI *train* function^54^ (scvi-tools version 0.16.4, following the versioning in ref. 29. Training on the concatenation achieved a slightly better fit than training performed only on the query data, as gauged by visualizing the final latent dimensions on a UMAP embedding and observing the overlap between studies within the “gdT” cluster, though, again, similar results were identified in both instances. As expected, the query data almost exclusively mapped to the “gdT” reference cluster (1,011/1,334 cells), confirming that the query cells were transcriptionally similar to annotated γδ T cells from other datasets.

To focus on the γδ T cells, all cells from the query and reference data that were annotated in the “gdT” and “gdT_naive” clusters (14,345 and 2,586 cells, respectively, from the combination of query and reference), as gauged by a weighted *k*-NN classifier following the design in ref. 29 were selected for downstream analysis. While we considered filtering based on annotation probability, that probability had low values in all cells that presented at the border between two clusters, even though they expressed genes characteristic of the cell type in which they were classified. We reasoned that a filter on probability would therefore likely discard meaningful signal and proceeded without one.

However, to ensure gut-resident γδ T cells had a robust, consistent transcriptional signal, we subjected the clusters to a round of additional filtering that removed: (1) all cells that were collected from tissues other than the small and large intestine (1,941 cells); (2) cells from Leiden subclusters of the “gdT” and “gdT_naive” groups, which were calculated from the same 30-nearest-neighbor graph of the 20 scVI dimensions used for the UMAP embedding, where the subclusters did not highly express *TRG* or *TRD* genes and/or did highly express *TRA* or *TRB* genes, as compared to the other subclusters (1,837 cells); (3) cells from individuals with 10 cells or less in the “gdT” and “gdT_naive” groups (151 cells from 31 individuals); and (4) cells from a single study^55^ that were segregated apart from cells from the other studies in the UMAP embedding (4,386 cells).

These filtering steps resulted in 8,859 cells for downstream analysis. To visualize the data, a standard analysis pipeline was subsequently performed that once again calculated a UMAP embedding (e.g., Figure 4A) from a 30-nearest-neighbor graph of the 20 scVI dimensions. Gene expression was visualized (e.g., Figure 4B) after normalization using the *SCTransform* function in Seurat (version 5.2.1)^56^, with separate models fit for each study via Seurat’s data layers design.

### Topic modeling

Traditional probabilistic topic modeling has no integration step to account for inter-individual or inter-study variability^30,31,45^. To remove signal from inter-individual variability in the topic model, we used the scANVI-corrected counts as input, instead of the original counts matrix, where each cell was given a size factor equal to the median number of UMIs detected across all γδ T cells in the joint query-reference dataset. This counts matrix consisted of only highly variable genes, since the published scANVI model was trained using only these genes. Further, to be used in the topic model, genes were required to be expressed in at least 3 cells in the data, and mitochondrial, ribosomal, and TCR genes were discarded. This filtering had several aims: (1) focusing on effector programs independent of TCR gene segment usage, (2) avoiding learning topics heavily influenced by mitochondrial or ribosomal genes, and (3) focusing on genes expressed in at least a handful of cells. Altogether, this resulted in a scANVI-corrected counts matrix filtered for topic modeling with positive gene expression of 1,145 genes in the γδ T cells. It is possible that a higher-resolution analysis using a larger number of genes would have yielded additional insights; however, this would have required re-training the reference model, which we opted against.

Topic modeling was then performed using the fastTopics package in R (version 0.6-192)^45,57^ for each *k* (the number of topics) ranging from 2 to 10, and for five random seeds for each *k*. For each topic model fit, topic-modeling-based differential expression (DE) analysis was performed on the scANVI-corrected counts using fastTopics, as detailed in the next section^58^. Topic model fits were evaluated based on a variety of factors, including the conservation of topics between studies in the dataset; the conservation of fits across the five random seeds; and how well the topic weights characterized gene expression profiles identified with the DE. Using these criteria, we selected *k*=3 for the number of topics, as larger *k* yielded programs less well conserved across publications, and *k*=2 merged the hybrid cytotoxic/NK-like topic (Figure 4A, Topic 3) with the naïve-like topic (Figure 4A, Topic 1).

### Differential expression analysis

We performed two types of DE analysis to evaluate genes in a topic. First, as part of the topic modeling pipeline applied to scANVI-corrected counts, we used the *de_analysis* function from the R package fastTopics^58^. This single-cell DE method incorporates the real-valued topic weights into a Poisson-based DE model to evaluate whether genes are enriched in any topic. It has advantages over using the raw gene weights calculated in the topic model fit, which do not include an estimation of statistical confidence and do not account for the differences between genes in their ranges of expression (e.g., *IFNG* is more lowly expressed in the data than *CD69*, but both are clearly enriched in topic 2). For our DE analysis, the “vsnull” approach was used to evaluate whether genes are DE in any topic compared to a null model consisting of a single topic for the entire data.

Second, to explicitly account for inter-individual variation in the statistical DE model, we developed a topic-based pseudo-bulk DE analysis approach that was applied to the original counts matrix. Specifically, for each topic, we separated cells into two bins: those that had “low” or “high” topic weights (Figure S6E, Table S4). These thresholds were selected based on the observed distributions of the topic weights both in the overall data and specifically in the data collected for this study, with the aim of dividing observed peaks located near 0 (ideally designated as “low” cells) from cells expressing a meaningful amount of the topic. Then we ran a traditional pseudo-bulk DE analysis, wherein gene expression counts for cells from each combination of topic bin and individual were aggregated into a single value, resulting in a pseudo-bulk gene expression matrix. We then calculated DE genes between the “high” and “low” bins using the limma-voom pipeline (version 3.58.1)^59,60^ with formula *∼0 + topic_bin*.

Individual-specific effects were modeled as a random effect using limma’s *duplicateCorrelation* function. To be included in the DE analysis for a specific topic, an individual was required to have at least 3 cells in both the “high” and “low” bins for that topic. Approaches with higher thresholds (e.g., 5 or 10 cells) more common in pseudo-bulk analysis were also implemented. However, these approaches discarded larger fractions of the cohort and resulted in more DEGs, suggesting that the lower threshold may be more conservative. Hence, we proceeded with the lower threshold for analysis. To further account for low cell number, the function *voomLmFit* was used to better account for zeros in the counts matrix, as opposed to the traditional *voom*^61^.

All genes presented in the figures were required to be significantly enriched in a topic according to both DE methods, provided the gene was included in the scANVI highly variable genes. Otherwise, if the gene was not in the scANVI model, only significance under the pseudo-bulk method was required. This same criterion was applied to determine the topic gene modules (Figures 4E–G, 4J–K, S6H-K, and S7E–F).

### Gene set enrichment analysis

Rank-based gene set enrichment analysis (GSEA) was performed using the *fgsea* package (version 1.28.0)^62^ in R. Genes were ranked based on the test statistic *t* from the topic-based pseudo-bulk differential expression. Enrichment analysis was performed using three databases: Gene ontology biological processes (GO BP)^63^, the Kyoto Encyclopedia of Genes and Genomes (KEGG)^64^, and the hallmark Molecular Signatures Database (MSigDB) gene sets^65^. After enrichment testing, pathways were collapsed to reduce the number pathways with strongly overlapping leading edges, using the *fgsea collapsePathways* function with a *pval.threshold* parameter setting of 0.1, though some overlaps still persisted. Analyses with all three processes are presented in Table S6. Curated GO BP pathways were selected for presentation in Figure 4D based on biological interpretability, with a preference for pathways whose genes did not strongly overlap with other selected pathways.

### Gene module scoring

Prior to scoring, an additional scANVI processing step was applied to filter out αβ T cells from Kornberg *et al.* and our frozen batch, which was spiked with αβIELs and included the majority of TCR clones with tested BTNL3/8 reactivity (Table S1). Specifically, a scANVI model was trained that included frozen cells (i.e., our frozen batch and Kornberg). Then, only cells from the batches which also included αβ T cells that had predicted “gdT” or “gdT_naive” annotations from the weighted kNN classifier were selected for downstream analysis. No additional filtering was performed on the batches from our publication that were not spiked with αβIELs, as these cells had already been determined to be TCRγδ^+^ from flow cytometry gating.

Scores for the three topics (Figures 4E, S6H, S6J, S7E–F) were then calculated from DE genes based on adjusted p-value threshold of <0.05 and a log-fold change threshold of >1 in absolute value in the relevant pseudo-bulk DE analyses (Table S5). We verified that trends persisted across choices of these thresholds. All genes included in the scANVI model and topic modelling were additionally required to have a fastTopics single-cell DE local false sign rate of <0.05 and posterior log-fold change estimate >0.5.

The core NK-like module scoring (Figures 4L–M, S7C–D, and S7G) used the genes *FCER1G*, *TYROBP*, *SH2D1B*, *FES*, and *RHOC*, which were isolated from the larger Topic 3 (hybrid cytotoxic/NK-like) module via directed comparisons to public data (Figures 4F–G and 4J–K). For random sampling (Figure S7G), subsets consisting of 5 genes were selected from the Topic 3 (hybrid cytotoxic/NK-like) module (Table S7).

Several module scoring methods have been proposed for single-cell^66–68^ and bulk^69–72^ RNA-seq data. Due to technical differences in single-cell versus bulk RNA-seq data, particularly that of dropout, preferred module scoring methods differ between the two modalities^68,73^. We accordingly used different scoring methods for analyses that scored clones (Figures 4L, S6J, S7C–E), which sometimes consist of only one cell, compared to those that used pseudo-bulk counts from various samples (Figures 4E, 4M, S6H, and S7F–G).

For clones, we used Seurat’s *AddModuleScore* function^56,66^, due to its frequent use in the field. Specifically, the scRNA-seq data was first normalized using the *SCTransform* function from Seurat, with separate models fit for each sequencing batch via Seurat’s data layers design. The gene module score was calculated using Seurat’s *AddModuleScore* on a cell-by-cell basis. To then link a module score to relative percent TCR downregulation (Figures 4L, S6H, S7C–E, and S7G), the data was subset to cells with TCRs whose BTNL3/8 reactivity had been tested, and the score was averaged across cells within each clone (i.e., with the same TCR), resulting in one score per clone. A single clone with tested reactivity (IEL TCR1/clonotype102_10X1 in Table S2) from the frozen batch was excluded because it was annotated as a “Trm_CD8” by the kNN classifier.

For pseudo-bulk module scores, we used the well-established approach of gene set variation analysis (GSVA) scores, calculated via the R GSVA package (version 1.50.0)^69^, which scales genes based on their expression level to ensure that more highly expressed genes are not more strongly weighted in the scoring. In the context of evaluating differences between haplotypes (Figures 4M and S7F–G), a pseudo-bulk matrix was calculated from the healthy control samples from this publication and Kornberg *et al.*^14^, filtered to remove lowly expressed genes using edgeR’s *filterByExpr*, (version 4.0.16) and normalized to LCPM values using limma’s *voom* function. The initial gene selection criterion for topics (Figure S7F), the core NK-like module (Figure 4M), and random Topic 3 (hybrid cytotoxic/NK-like) genes (Figure S7G) was the same as for the clone-based scoring, but any genes in the modules that did not pass filtering for the combined McDonald and Kornberg matrix were excluded from scoring.

For both types of scoring, we also found consistent trends across several methods (clone-based scoring: Seurat’s *AddModuleScore*, AUCell, Jasmine, average expression, and average scaled expression; pseudo-bulk scoring: GSVA, ssGSEA, singscore, z-scoring, and average expression)^66–72^.

### Use of public MMRd CRC and celiac disease scRNA-seq datasets

There exist several public scRNA-seq datasets of intestinal diseases. The two disease datasets used in this publication^37,38^ were chosen in part because the large numbers of individuals profiled in the two cohorts adequately powered the pseudo-bulk DE analysis. Cell types in each dataset were annotated using the same scANVI gut reference atlas as applied to our samples. To isolate CD8 T cells from MMRd CRC, cells annotated as “Trm_CD8”, “Trm/em_CD8”, and “Tnaive/cm_CD8” were selected. To isolating γδ T cells from both datasets, the same pipeline was applied as for our healthy query-reference atlas; i.e., all cells annotated as “gdT” or “gdT_naive” were selected, and then an additional round of filtering was performed to remove clusters that did not highly express *TRG* or *TRD* genes, or that did highly express *TRA* or *TRB* genes. We considered an analogous filtering step for the CD8 T cells, but TCR gene expression was relatively homogeneous among these cells, with detectable *TRA* and *TRB* expression, so no additional filtering was applied. To examine epithelial cell expression of *BTNL3* and *BTNL8*, cell type annotations from the original publications were used.

For DE analysis, a pseudo-bulk pipeline comparing disease to healthy samples in each publication was applied using *limma-voom*. To be included in these comparisons, an individual was required to have at least five cells of the cell type of interest. As before, the function *voomLmFit* was used to better fit low count data. Due to low power in the γδ T cells, likely because of low cell numbers, the adjusted P-value threshold was increased to <0.1 (from <0.05). Moreover, a slightly lower log-fold change threshold of >0.5 (rather than >1) in absolute value was used in the disease context DE analyses, motivated in part by the identification of several Topic 3 (hybrid cytotoxic/NK-like) genes that were significantly DE in disease, but with a slightly lower log-fold change than in the healthy context.

To estimate the statistical significance of overlap between gene sets (Figures 4J and S6K), gene set overlaps were compared to the number of overlapping genes from 10,000 simulations of an estimated null distribution, consisting of three independent binomial distributions describing the genes DE in each dataset, calculated using *rbinom* in R. For each comparison, we used the fraction of genes overlapping between the three observed DE gene lists as the draw probability for the binomial distribution, while the total number of draws was equal to the total number of genes expressed in all three datasets. P-values were then calculated by determining the fraction of empirical null simulations in which the number of overlapping genes was greater than or equal to the observed number of overlapping genes.

### Stimulation and Library Preparation for bulk RNA-seq on ex vivo CD8 IELs

IELs collected from six individuals were either left unstimulated or were stimulated for 3h with αCD3 (BD; clone UCHT1; 1.5μg/mL) in flat-bottomed 96-well plates (Corning; 3370). Following stimulation, the cells were stained and 500 live, CD45^+^ CD8β^+^ CD4^-^ cells were sorted into semi-skirted LoBind twin.tec PCR plates (Eppendorf) containing 12μL sorting solution, as described in the SMART-Seq v4 Ultra Low Input RNA Kit and flash-frozen at -80°C until all replicates were collected. CD3 and TCRαβ were not used for gating as they are downregulated following αCD3 stimulation. cDNA was then generated from the sorted cells according to the manufacturer’s specifications.

RNA-seq libraries were generated using 200 pg of purified cDNA using the Nextera XT DNA Library preparation kit (Illumina) and IDT for Illumina Nextera DNA Unique Dual Indexes (Illumina). Obtained libraries were quantified using the Agilent High Sensistivity DNA Kit (Agilent), normalized, multiplexed, and sequenced at a depth of 20 million reads per sample (single read 100bp) on a NovaSeq X (Illumina) at the University of Chicago Genomics Facility.

### Bulk RNA-seq preprocessing

Data was processed using the nf-core/rnaseq (v3.14.0) pipeline^74^. Briefly, raw reads were trimmed using TrimGalore! (v0.6.7), mapped to the Human GRCh38 reference genome using STAR (v2.7.9a)^75^, and quantified using Salmon (v1.4.0)^76^ and gene annotation data from Gencode V34 to produce the raw count matrix.

### Bulk RNA-seq differential expression analysis

To compare CD8 IELs that were stimulated with αCD3 to those that were left unstimulated, we applied a limma-voom differential expression pipeline similar to that used for pseudo-bulk DE. Specifically, genes with low counts in any experimental group were filtered out using edgeR’s *filterByExpr* function, with the minimum count required to pass filtering set to the default value of 10. Then, library sizes were scaled via edgeR’s *normLibSizes* function, using the trimmed mean of M-values (TMM) method. A principal component analysis was performed on the log CPM counts, which identified strong transcriptional effects associated with the individual from which the cells were sampled. Therefore, for the differential expression analysis, we chose to use a design formula which explicitly modeled individual: *∼ 0 + stimulation + individual*, from which we fit a contrast comparing the stimulated and unstimulated coefficients. Modeling individual as a random effect, as in the pseudo-bulk DE, produced comparable effects and resulted in identification of the same set of core NK-like genes.

## Notes

### Competing Interest Statement

The authors have declared no competing interest.

