## Supplementary Figures for "Adaptive-like features of the γδ TCR couple chronic BTNL recognition to NK-like tissue immunity"

Figure S1. TCRs selected for BTNL3/8 reactivity assays are representative of the observed Vγ4 TCR repertoire in their respective tissue compartments, related to Figure 1.

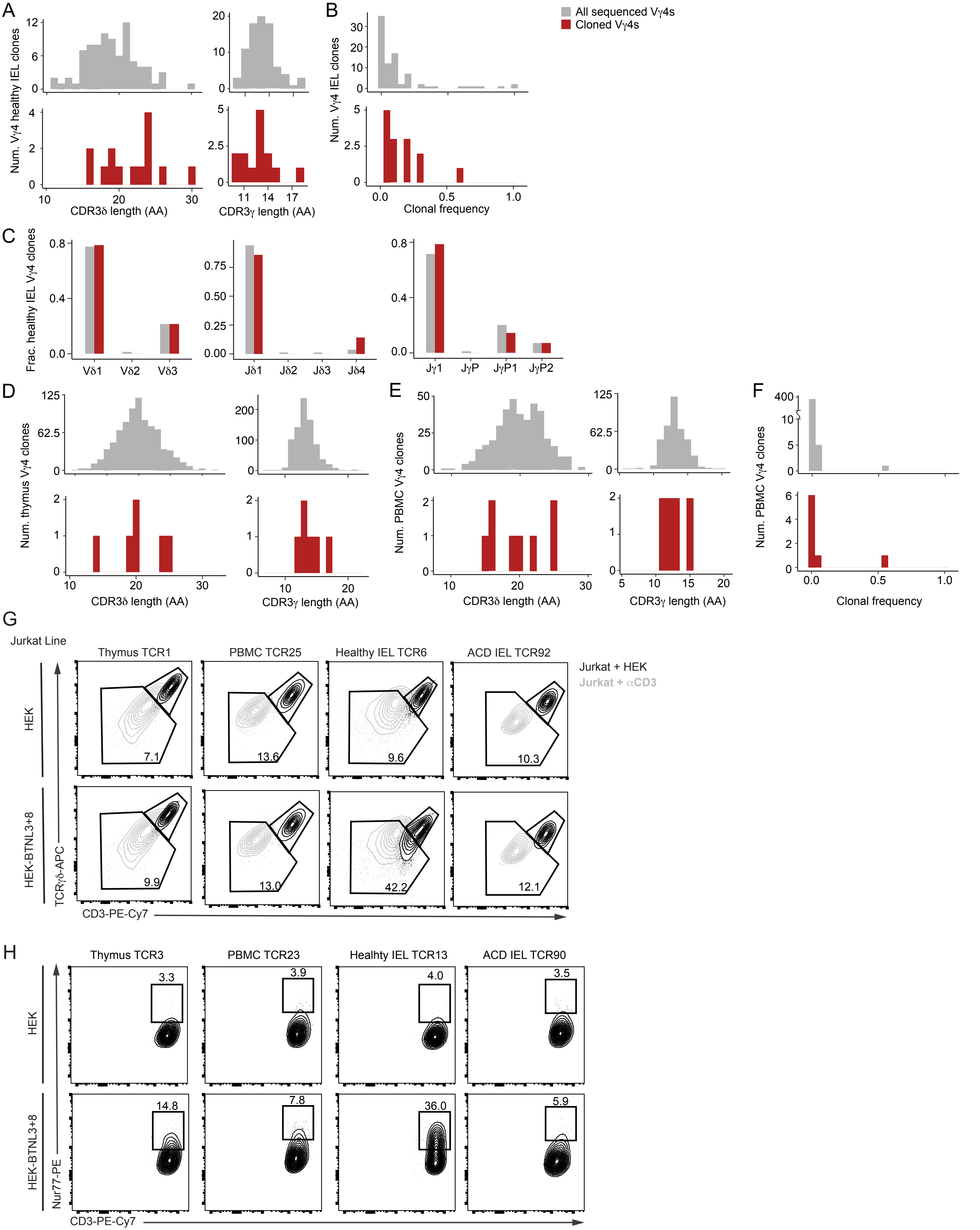

Figure S2. Haplotypic variants of *TRGV4*, consisting of variable amino acids within and outside of the HV4γ region, show reduced BTNL3/8 reactivity, related to Figure 1.

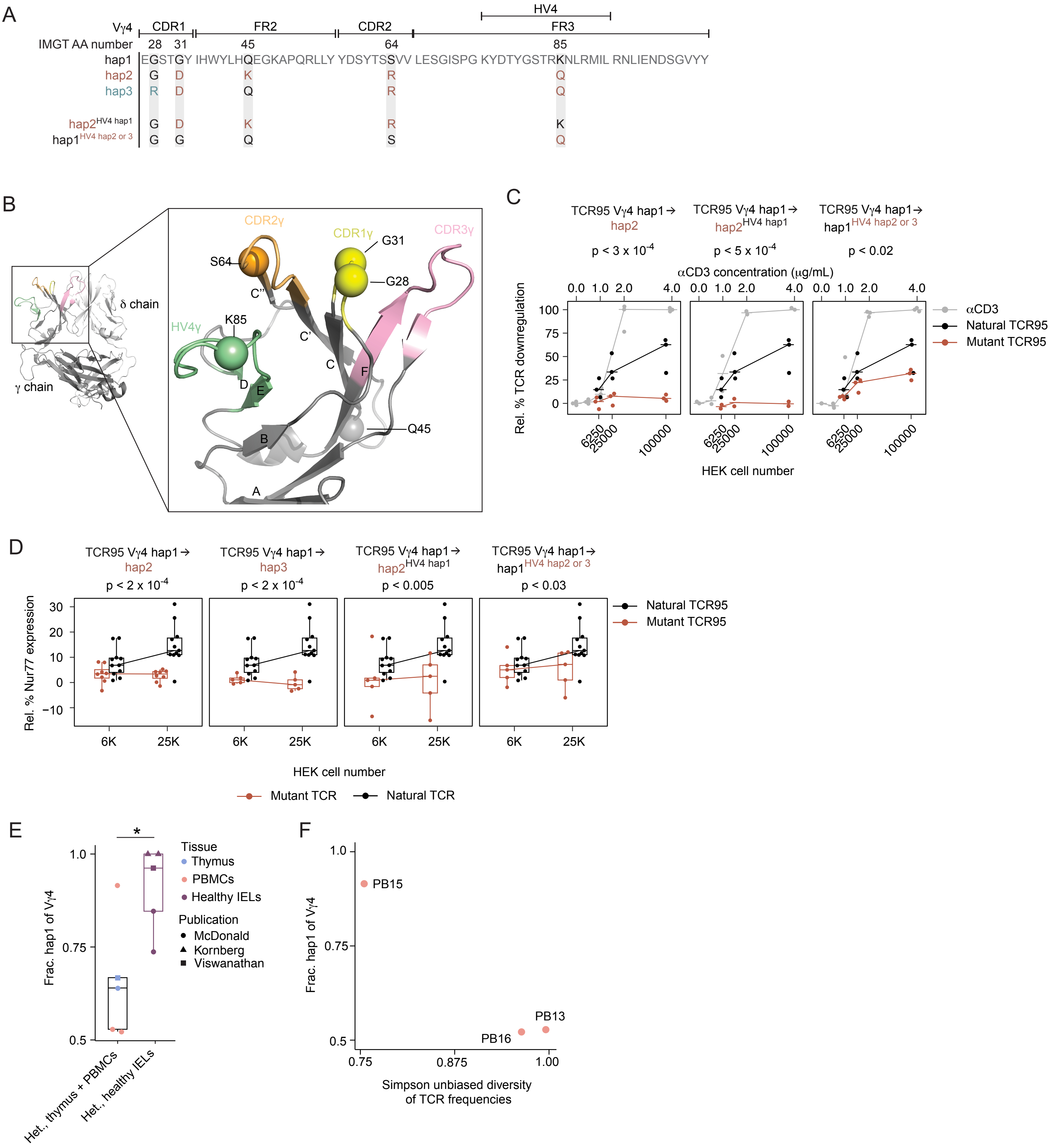

Figure S3. Specific γδ TCR traits are associated with tissue residence and reactivity to BTNL3/8, related to Figure 2.

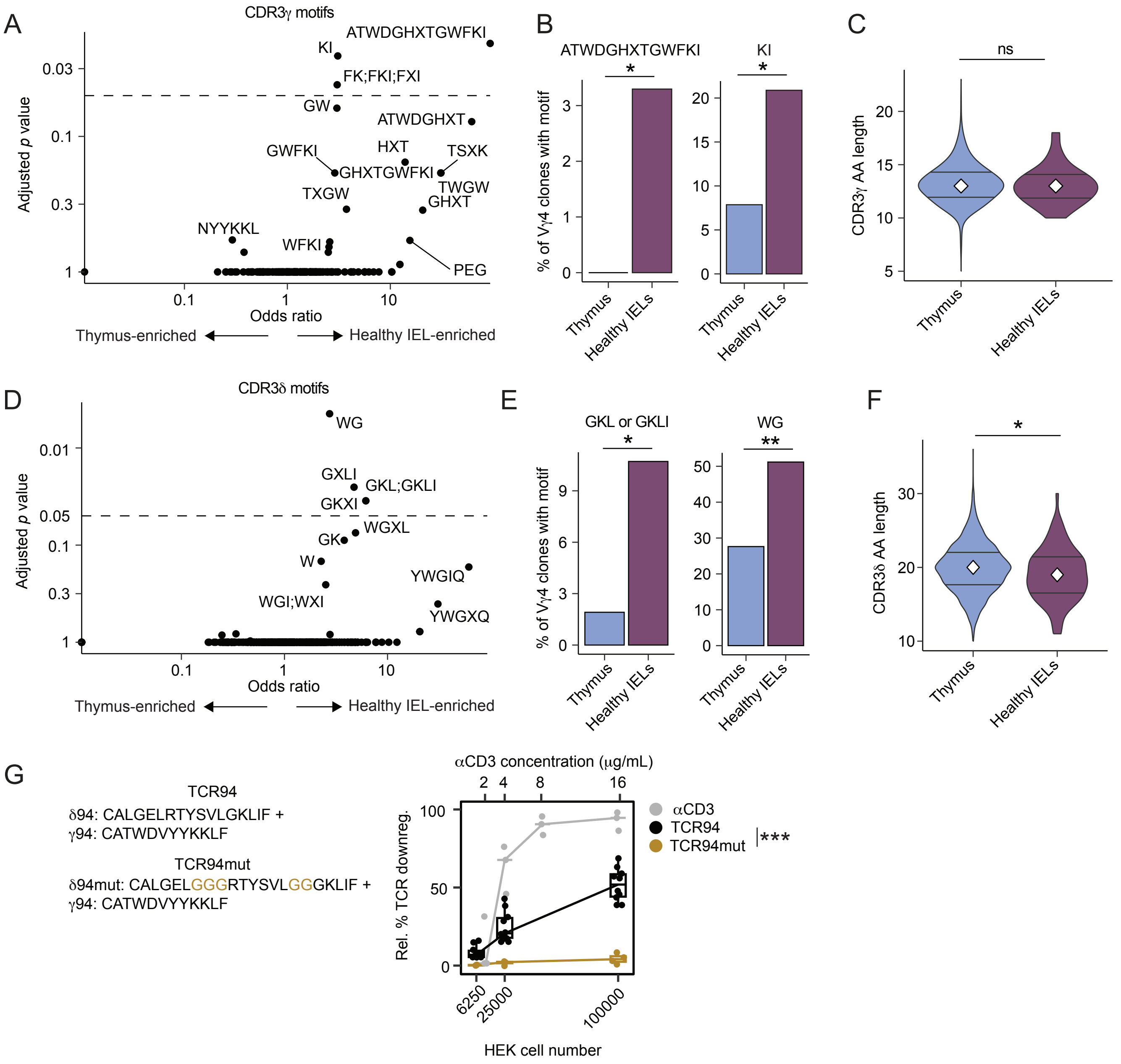

Figure S4. Haplotypic variants of *TRGV4* show comparable binding to BTNL3, related to Figure 3.

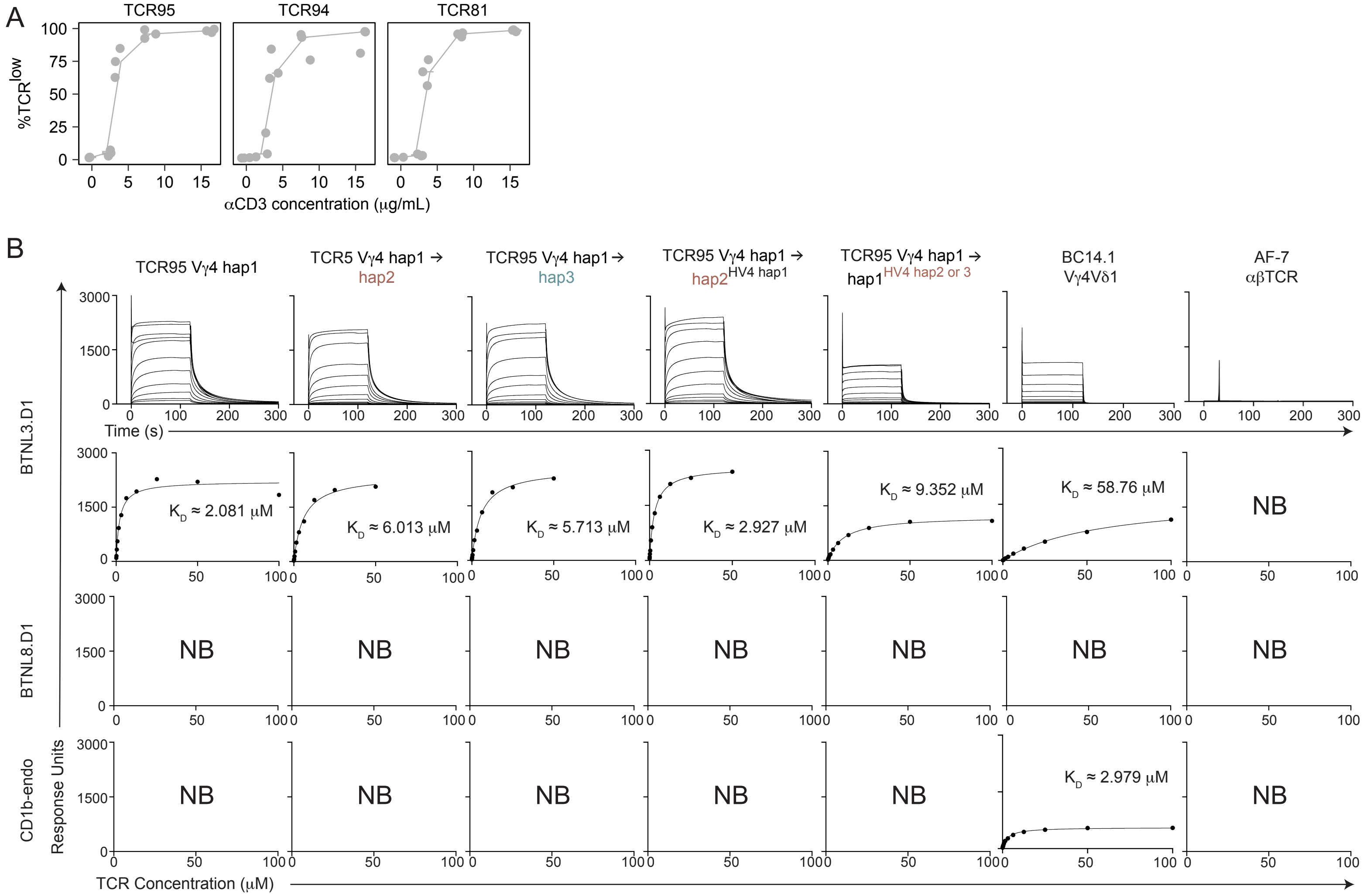

Figure S5. Haplotypic variants of *TRGV4* show partial ERK induction but no CD3 clustering or CD3ζ phosphorylation in response to BTNL3/8, related to Figure 3.

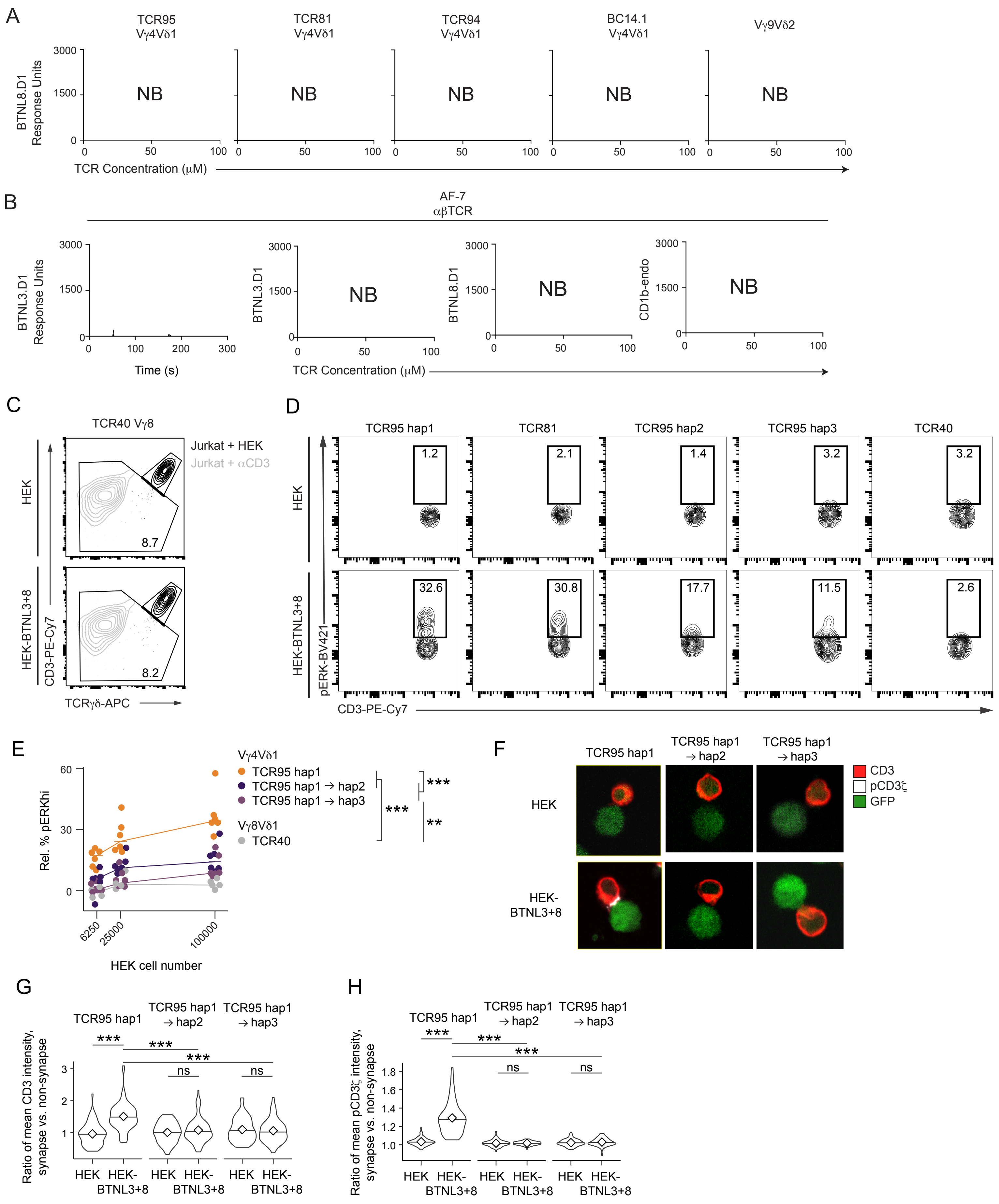

### Figure S6. Topic modeling identifies a gene program conserved between individuals and linked to BTNL3/8 reactivity, related to Figure 4.

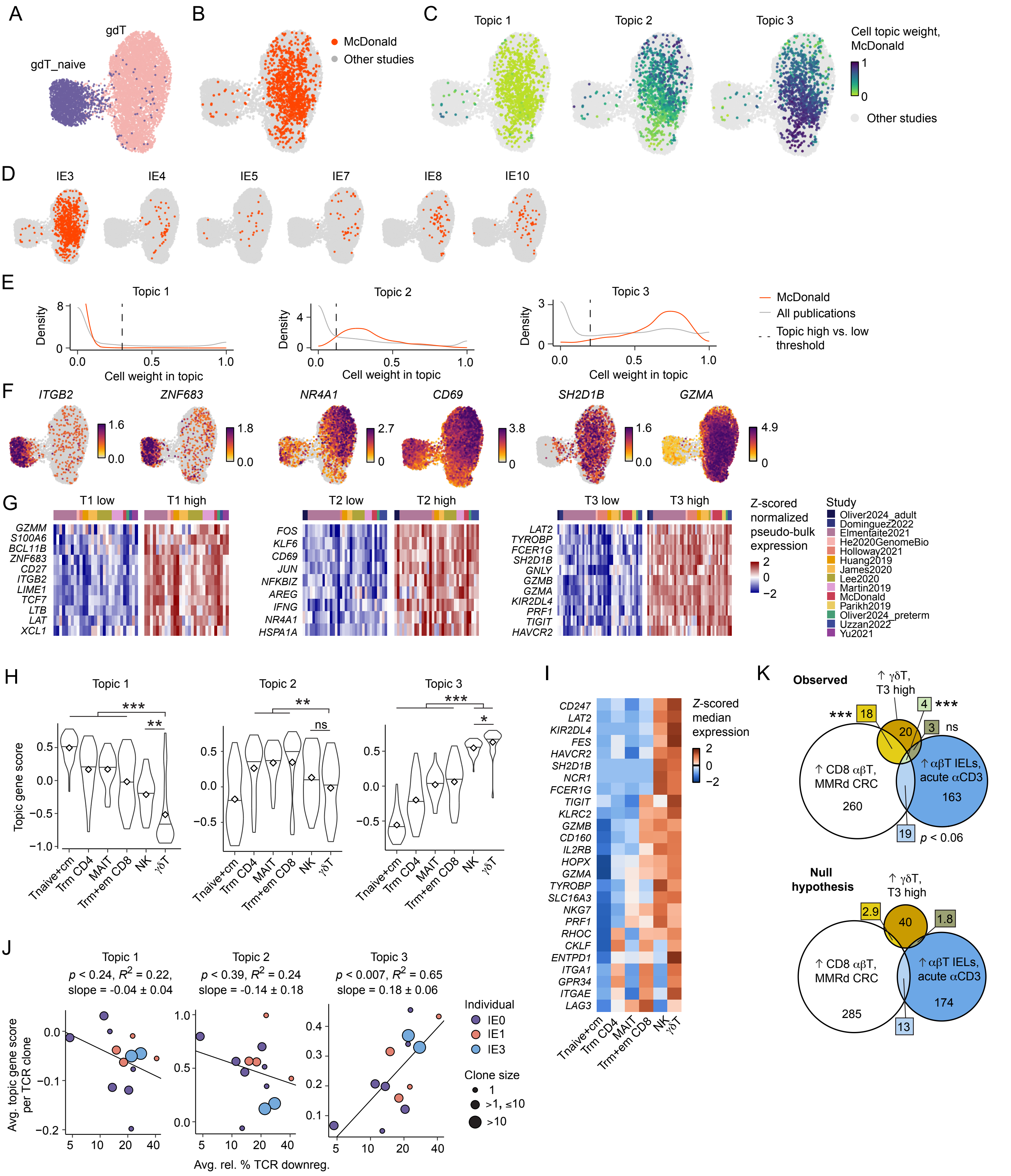

Figure S7. Topic modeling identifies an NK-like gene program characteristic of chronic antigen exposure, related to Figure 4.

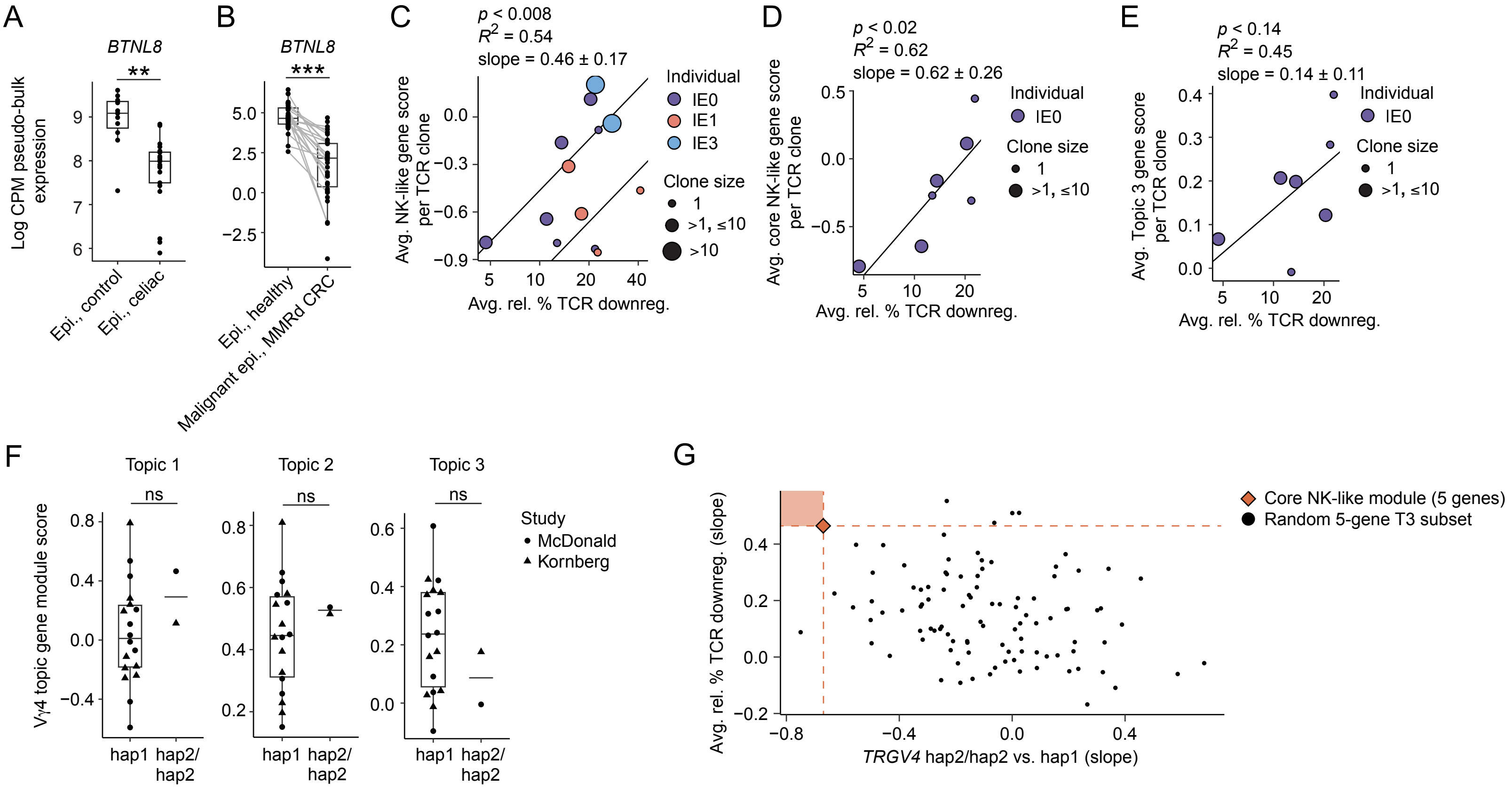
